# Lysine Demethylase 4A is a Centrosome Associated Protein Required for Centrosome Integrity and Genomic Stability

**DOI:** 10.1101/2024.02.20.581246

**Authors:** Pratim Chowdhury, Xiaoli Wang, Julia F. Love, Sofia Vargas-Hernandez, Yuya Nakatani, Sandra L. Grimm, Dereck Mezquita, Frank M. Mason, Elisabeth D. Martinez, Cristian Coarfa, Cheryl L. Walker, Anna-Karin Gustavsson, Ruhee Dere

## Abstract

Centrosomes play a fundamental role in nucleating and organizing microtubules in the cell and are vital for faithful chromosome segregation and maintenance of genomic stability. Loss of structural or functional integrity of centrosomes causes genomic instability and is a driver of oncogenesis. The lysine demethylase 4A (KDM4A) is an epigenetic ‘eraser’ of chromatin methyl marks, which we show also localizes to the centrosome with single molecule resolution. We additionally discovered KDM4A demethylase enzymatic activity is required to maintain centrosome homeostasis, and is required for centrosome integrity, a new functionality unlinked to altered expression of genes regulating centrosome number. We find rather, that KDM4A interacts with both mother and daughter centriolar proteins to localize to the centrosome in all stages of mitosis. Loss of *KDM4A* results in supernumerary centrosomes and accrual of chromosome segregation errors including chromatin bridges and micronuclei, markers of genomic instability. In summary, these data highlight a novel role for an epigenetic ‘eraser’ regulating centrosome integrity, mitotic fidelity, and genomic stability at the centrosome.

## INTRODUCTION

Centrosomes are evolutionarily conserved membraneless organelles with two orthogonally arranged mother and daughter centrioles surrounded by a cloud of pericentriolar material^1^. They provide a critical function during cell division organizing microtubules of the mitotic spindle and functioning as a MicroTubule Organizing Center (MTOC) at each pole of the spindle. Centrosomes duplicate once every cell cycle in S-phase, mature and acquire pericentriolar material in G2, and finally separate at the onset of mitosis to enable formation of a bipolar spindle^2^. The formation of the spindle is determined by the integrity of the centrosomes themselves, which ultimately ensures accurate chromosome segregation to the two ensuing daughter cells. The precise interactions that occur in space and time to orchestrate these centrosome specific events are key to maintaining mitotic fidelity^3^.

Loss of centrosome integrity results in structural, numerical, and/or functional aberrations associated with many diseases including cancer^4^. Structural perturbations can range from changes in centriole length to alterations in the amount and composition of the pericentriolar material^5^. Numerical aberrations, most associated with centrosome amplification, can arise from overduplication, fragmentation, or tetraploidization. Irrespective of the mechanism, centrosome amplification is linked to numerous pathological diseases and is a hallmark of most cancers^6^. An increase in centrosome numbers is linked to defective mitosis and chromosome segregation defects that likely contribute to acquisition of chromosome abnormalities and genomic instability^4^. Cells with multi-polar spindles arise as a consequence of having multiple centrosomes and these cells are usually eliminated via apoptosis. To avoid cell death, while advantaging genomic instability as an oncogenic driver, cancer cells develop strategies to avert mitotic catastrophe in the setting of centrosome amplification. Clustering centrosomes to assemble pseudo-bipolar spindles is one such strategy used by cancer cells to overcome the challenge of harboring multiple centrosomes^7^. This strategy allows successful progression through mitosis, although aberrant spindle assembly, damaged kinetochore attachments, and chromosome segregation errors that can ultimately cause aneuploidy and drive oncogenesis are the resulting payment^8^.

Lysine demethylase 4A (KDM4A) is a demethylase that belongs to the Jumonji family of enzymes that remove/erase methyl marks on histones^9^, with well-established catalytic and scaffolding functions on chromatin^9^. KDM4A demethylates i.e., ‘erases’ di- and tri-methylation on histone H3 lysine 9 and 36 residues (H3K9me2/3 and H3K36me2/3)^10, 11^. Our studies revealed that KDM4A also localizes to the centrosome, both in interphase and mitotic cells where it interacts, and co-localizes, with centrosome-associated proteins. KDM4A maintains centrosome homeostasis with loss of *KDM4A* resulting in supernumerary centrosomes and disruption of normal mitosis. Defective mitosis in *KDM4A*-null cells results in the accrual of chromosome segregation errors including chromatin bridges and formation of micronuclei, a marker of genomic instability. KDM4A enzymatic function is required for maintaining centrosome integrity and genomic stability, although the role of KDM4A in regulating centrosome integrity appears unrelated to regulation of gene expression. In summary, our data identify a novel function for an epigenetic eraser in regulating centrosome integrity, mitotic fidelity, and genomic stability.

## RESULTS

### KDM4A is a centrosome associated protein

We found, in addition to its known localization in the nucleus, robust cytoplasmic localization of KDM4A in human (HEK293T and human kidney cells (HKC)) **(Figs. S1A-B)** and mouse (MEFs) cells **(Fig. S1C)** using subcellular fractionation assays. This dual localization (nuclear and cytoplasmic) was validated with antibody-independent assays using exogenously expressed hemagglutinin-tagged HA-KDM4A in human cells (HEK293T and HKC) **(Figs. S1D-E)**. Spatial localization of KDM4A in cells, was assessed using direct epifluorescence microscopy, which revealed that in addition to the nuclear localization during interphase, KDM4A also displayed strong focal immunoreactivity at centrosomes of interphase cells **(Fig. 1A, and Figs. S1F-G)**, and at the spindle poles in cells during mitosis **(Fig. 1A)**. We also followed KDM4A at the centrosomes across mitosis in hTERT immortalized RPE-1 (retinal pigmented epithelial) cells. As shown in **Figs. 1B-C**, KDM4A immunoreactivity was detected at the spindle poles from prophase through telophase, where it co-localized with two centrosome-specific markers: gamma (γ)-tubulin **(Fig. 1B)** and GFP-centrin **(Fig. 1C)**, which was further validated using an antibody-independent approach with RFP-tagged KDM4A, which confirmed spindle pole localization during mitosis **(Fig. 1D)**.

**Figure 1.**
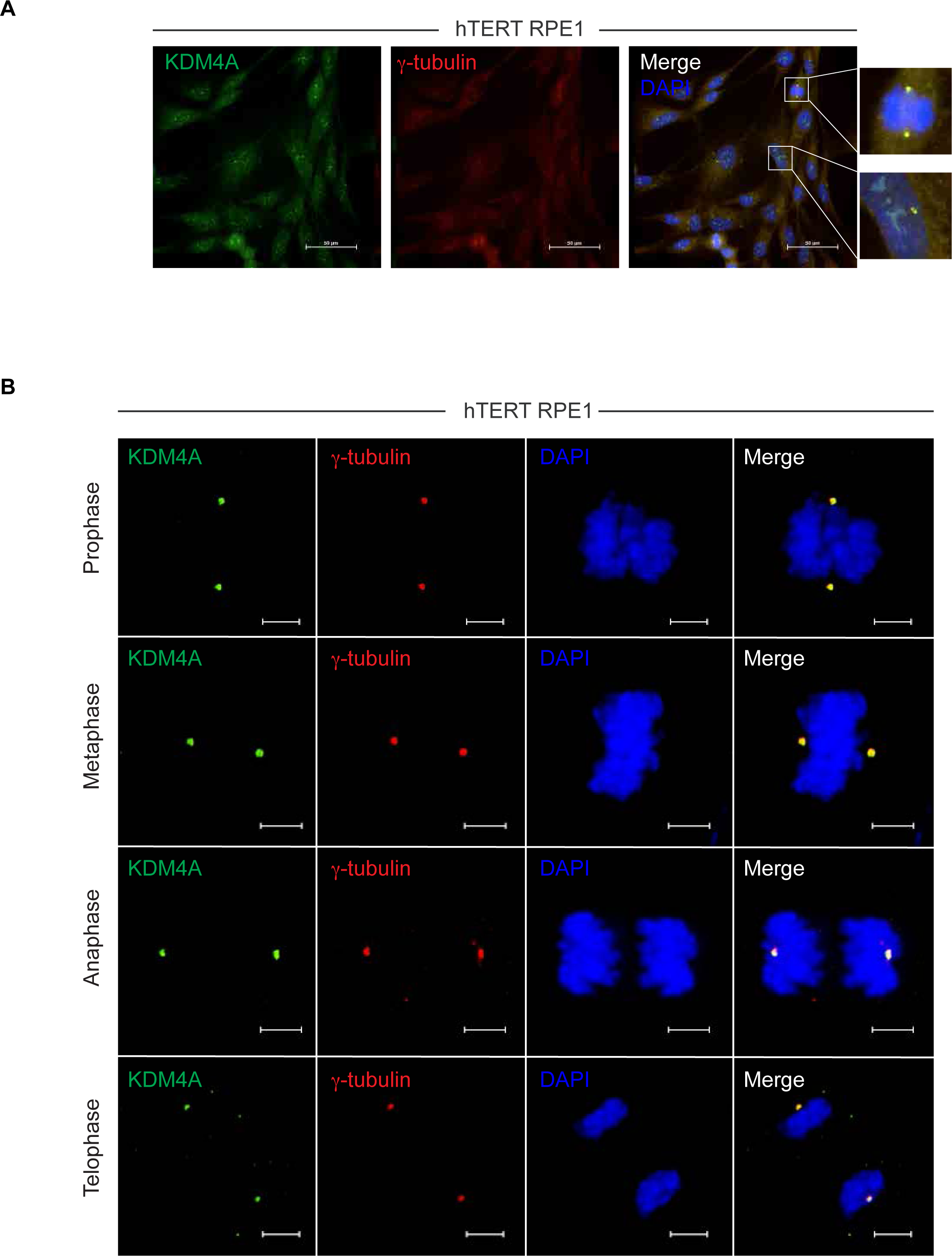
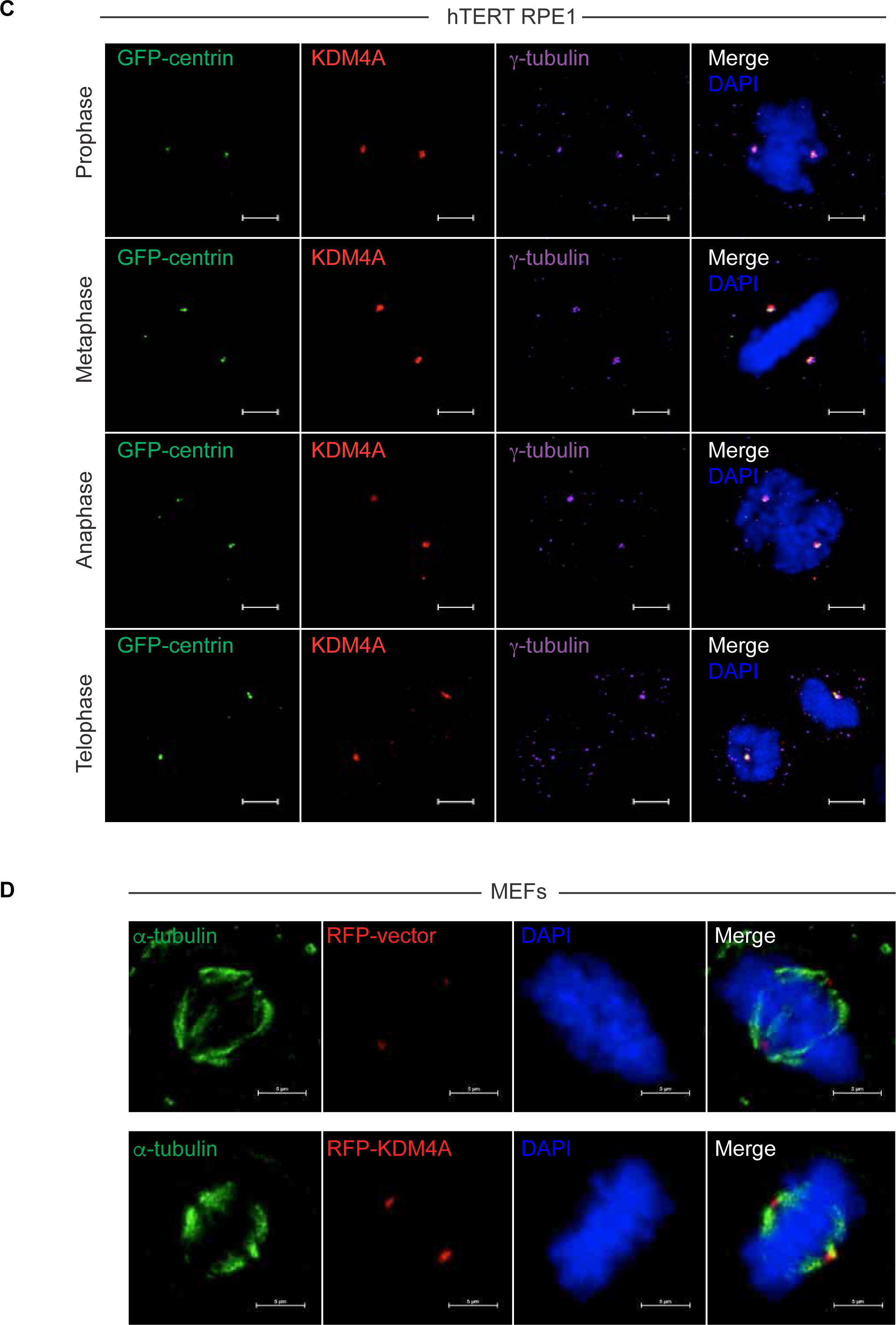
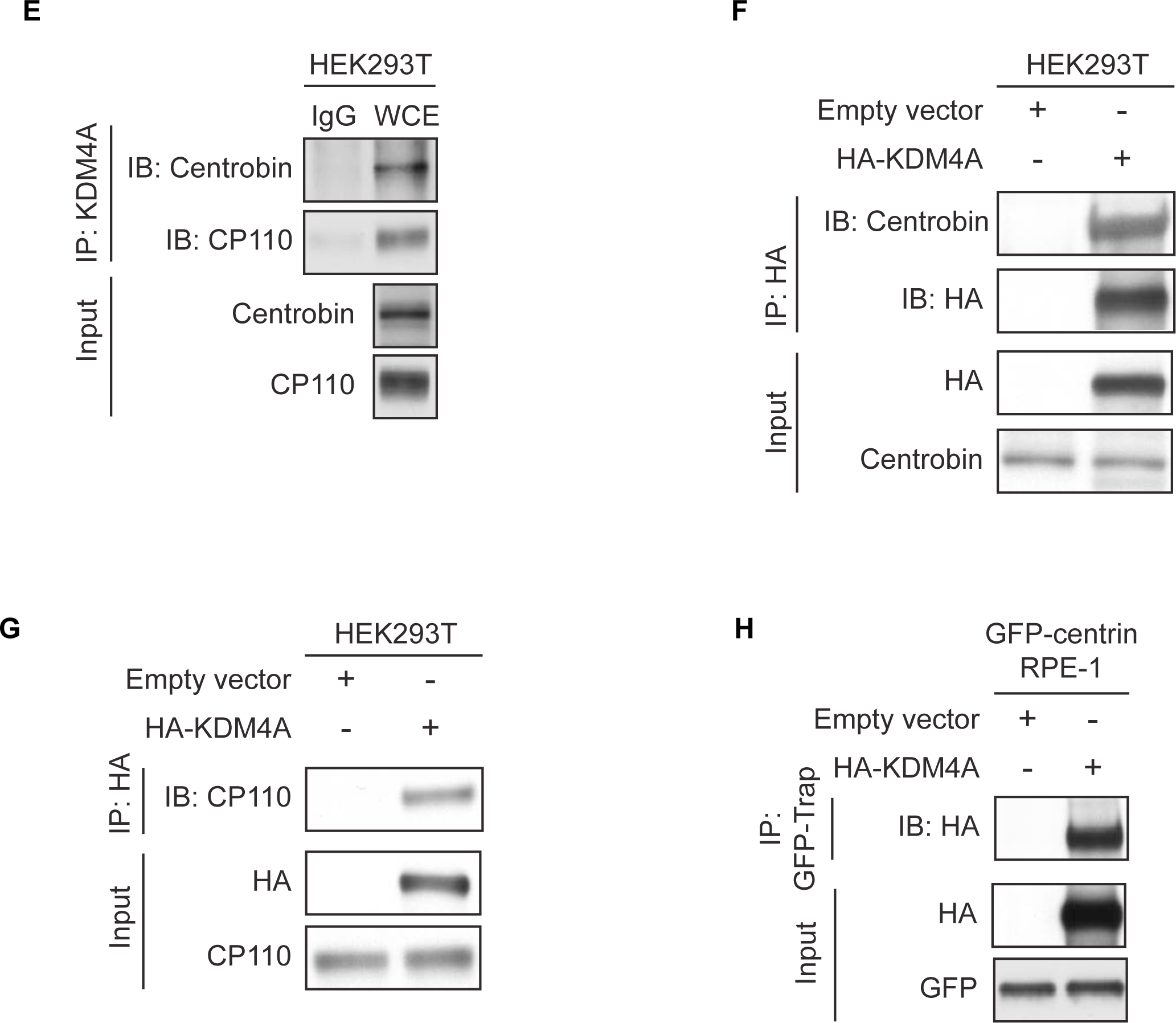
KDM4A is a centrosome-associated protein. A Representative immunofluorescence image showing localization of KDM4A (green) in hTERT RPE-1 cells using anti-KDM4A antibody. The cells were labelled for α-tubulin (microtubule marker, red) and counterstained with DAPI (nuclei, blue). Scale bar – 50 μm. B Representative images showing KDM4A staining (green) in the four phases of mitosis (prophase, metaphase, anaphase, and telophase) in hTERT RPE-1 cells. These cells were labelled for α-tubulin (microtubule marker, red) and counterstained with DAPI (nuclei, blue). Scale bar – 5 μm. C Representative images of the four phases of mitosis in hTERT RPE-1 cells stably expressing GFP-centrin (centrosome marker, green). These cells were labelled for KDM4A (red) and γ-tubulin (centrosome marker, violet) and counterstained with DAPI. Scale bar – 5 μm. D Representative images of mouse embryonic fibroblasts (MEFs) transiently expressing Lentizeo-RFP-vector (upper panel, red) or Lentizeo-RFP-KDM4A (lower panel, red). The microtubules were labelled with α-tubulin (green) and the nuclei counterstained with DAPI. Scale bar – 5 μm. E Immunoblots showing co-immunoprecipitation of endogenous KDM4A with centrobin and CP110 (centrosome proteins) from HEK293T cellular extracts. Input lysates are also indicated as shown. F, G Immunoblots showing co-immunoprecipitation of exogenously expressed HA-KDM4A with centrobin (F) and CP110 (G) from HEK293T cellular extracts. Input lysates are indicated in each panel as shown. H Immunoblots showing co-immunoprecipitation of GFP-centrin with exogenously expressed HA-KDM4A. Expression of GFP-centrin and HA-KDM4A are indicated to show input samples.

In parallel, biochemical assays to probe potential interactions of KDM4A with centrosome-specific proteins were performed. Co-immunoprecipitation confirmed the ability of endogenous and exogenously expressed HA-KDM4A to bind centrobin **(Figs. 1E and 1F)** and CP110 **(Figs. 1E and 1G)**, respectively – two centrosome specific proteins that recognize mother (CP110) and daughter (CP110 and centrobin) centrioles. Reverse co-immunoprecipitation confirmed the association of KDM4A with both centrobin **(Fig. S1H)** and CP110 **(Fig. S1I)**. GFP-trap assays additionally confirmed the ability of GFP-centrin to also pulldown HA-KDM4A in hTERT RPE-1 cells stably expressing GFP-centrin **(Fig. 1H)**. These localization and biochemical data suggest that in addition to its known function as a chromatin remodeler, KDM4A may also have a functional association with the centrosome.

### Single-molecule localization microscopy validates KDM4A as a component of the MTOCs

To precisely position and quantify KDM4A, and further define the size and shape of centrosome-localized KDM4A, we used the single-molecule localization microscopy (SMLM) technique called exchange points accumulation for imaging in nanoscale topography (Exchange-PAINT)^12,13^ for two-target 3D super-resolution imaging. This antibody-based technology uses a short oligonucleotide strand, referred to as the ‘docking’ strand to label a secondary antibody. Multi-target imaging is performed by sequential flow-through of dye-labelled ‘imager’ oligonucleotide strands, which are complementary to the ‘docking’-strands. Transient binding of the ‘imager’-strands to the ‘docking’-strands generates apparent blinking that enables single-molecule localization. Our methodology employed Exchange-PAINT with microfluidics-assisted washes to achieve sequential imaging of two targets, together with point spread function (PSF) engineering to achieve 3D super-resolution imaging (**Fig. S2)** (see also Methods). Using this technology, 3D super-resolved reconstructions revealed a well-localized signal for KDM4A at the centrosomes, detected using the centriolar marker centrobin **(Figs. 2A-C and Figs. S3A-I)**, during mitosis analogous to our observations using diffraction-limited epifluorescence microscopy **(Fig. S3J)**. Importantly, the distribution of KDM4A extended outside of the centrobin distribution yielding diameters of 749 ± 79 nm (reported as mean ± standard deviation from 15 measurements in total of five KDM4A distributions) **(Fig. 2D and Figs. S4A-L, S4M-X, and S4AA-FF,** see also Methods). The quantification data of the diameter of the KDM4A distribution, independently in the different axes (x, y, z), revealed a relatively uniform/circular distribution **(Fig. 2D)** suggesting that KDM4A was not restrained to a particular region of the centrosome. These single-molecule data firmly establish KDM4A at the centrosomes of the spindle during mitosis.

**Figure 2.**
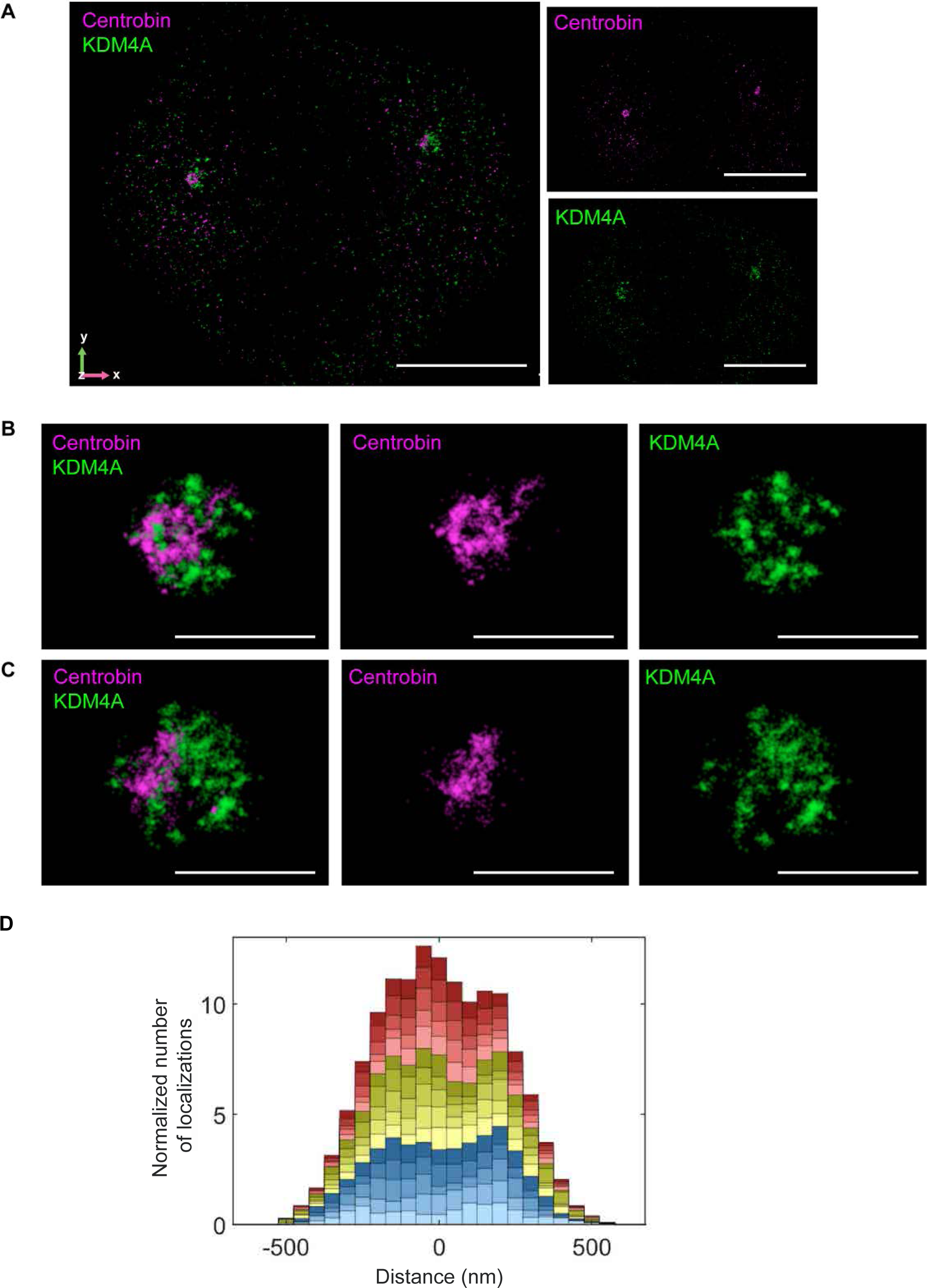
Single-molecule super-resolution reconstruction of KDM4A at the centrosomes during mitosis. A Representative 3D reconstruction showing the super-resolved distributions of KDM4A (pseudo-colored green) and centrosome marker – centrobin (pseudo-colored magenta) at the spindle poles during mitosis. The panels to the right show the centrobin data (top, pseudo-colored magenta) and the KDM4A data (bottom, pseudo-colored green) separately. Scale bar - 5 μm. B, C Magnified images of each of the spindle poles from (A) showing the KDM4A distribution (pseudo-colored green) overlapping with the distribution of the centrosome marker – centrobin (pseudo-colored magenta). The pseudo-colored individual reconstructions are shown to the right of the merged image. Scale bar – 1 μm. D Cumulative histogram of the normalized number of localizations for five KDM4A distributions along the x-direction (blue), y-direction (yellow) and z-direction (red). Data for the five individual distributions are shown by the color shading for each axis.

### KDM4A is required for the maintenance of centrosome number

Having established KDM4A as a centrosome-localized protein, we next investigated a functional role for KDM4A at mitotic centrosomes. *Kdm4a*-deficient MEFs were generated using CRISPR-Cas9 gene editing and *Kdm4a* knockout confirmed using immunoblotting for KDM4A protein levels in clonally derived cells and validated by increased histone H3 lysine 36 tri-methylation (H3K36me3) in *Kdm4a*-deficient cells **(Fig. 3A)**. To probe centrosome numbers in individual cells, we selected γ-tubulin as a marker for centrosomes in interphase cells, since cells normally contain a constant 2-γ-tubulin foci per cell. As shown in **Fig. 3B**, *Kdm4a*-null cells had increased γ-tubulin foci per cell (i.e., >2 foci per cell), consistent with centrosome amplification. Quantification revealed that *Kdm4a*-deficient cells (*Kdm4a*-CRISPR MEFs) had a significantly higher incidence of >2 γ-tubulin foci/cell compared to controls (non-targeting sgRNA MEFs) **(Fig. 3C)**. Increased number of γ-tubulin foci arise via multiple mechanisms, and to evaluate if the γ-tubulin foci in *Kdm4a*-null cells were comprised of centrioles, the *Kdm4a*-deficient cells were probed using anti-Cep135 (a centriole-specific marker) antibody. The supernumerary γ-tubulin foci in cells lacking *Kdm4a* stained positive for Cep135 **(Fig. 3D)** indicating centrioles in these foci. In addition to Cep135, these γ-tubulin foci were also positive for pericentrin **(Fig. 3E)**, a marker of the pericentriolar material that surrounds the centrioles. These data show KDM4A is required for maintenance of normal centrosome numbers in cells, and in the absence of this demethylase, results in supernumerary centrosomes.

**Figure 3.**
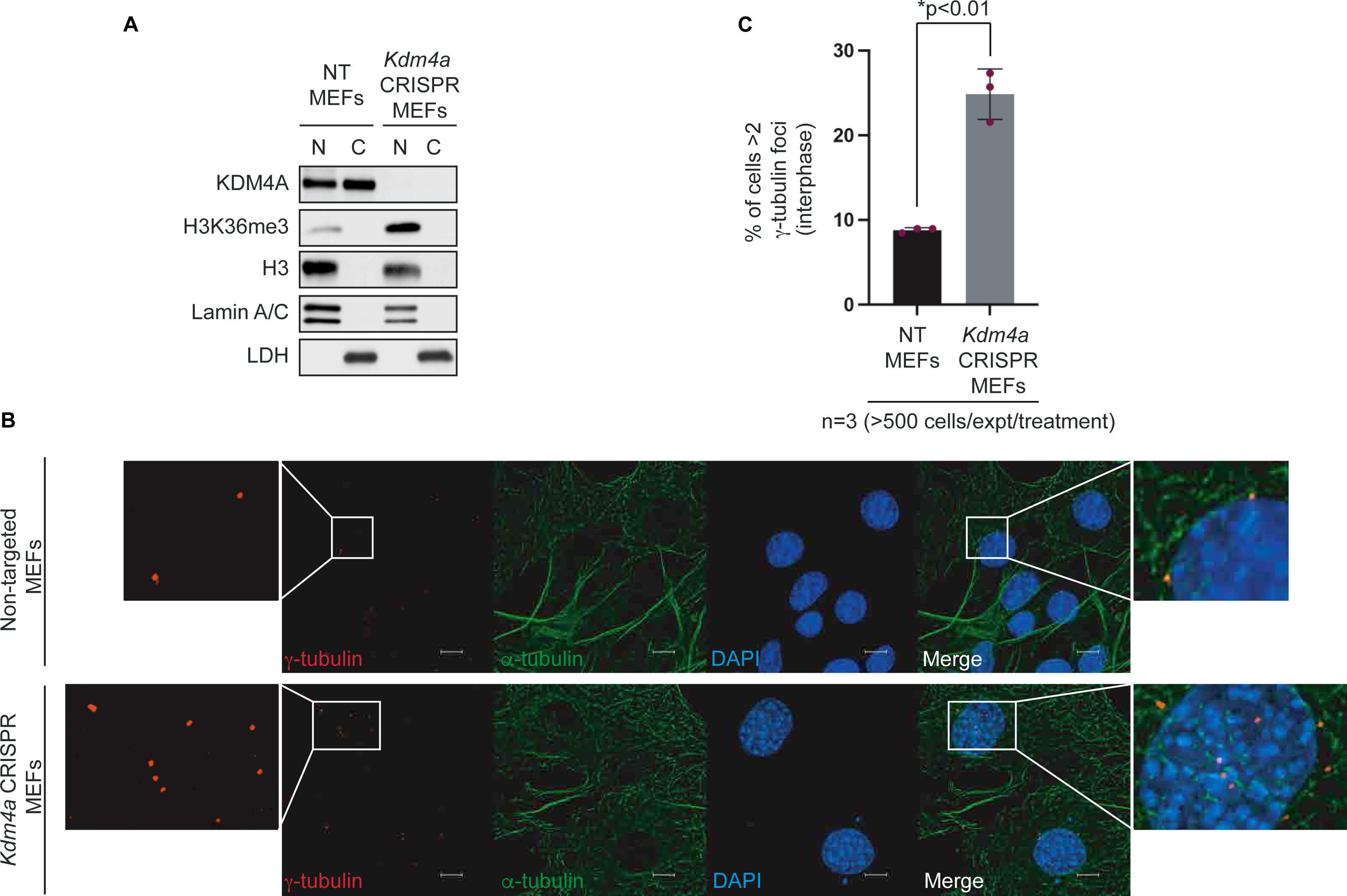
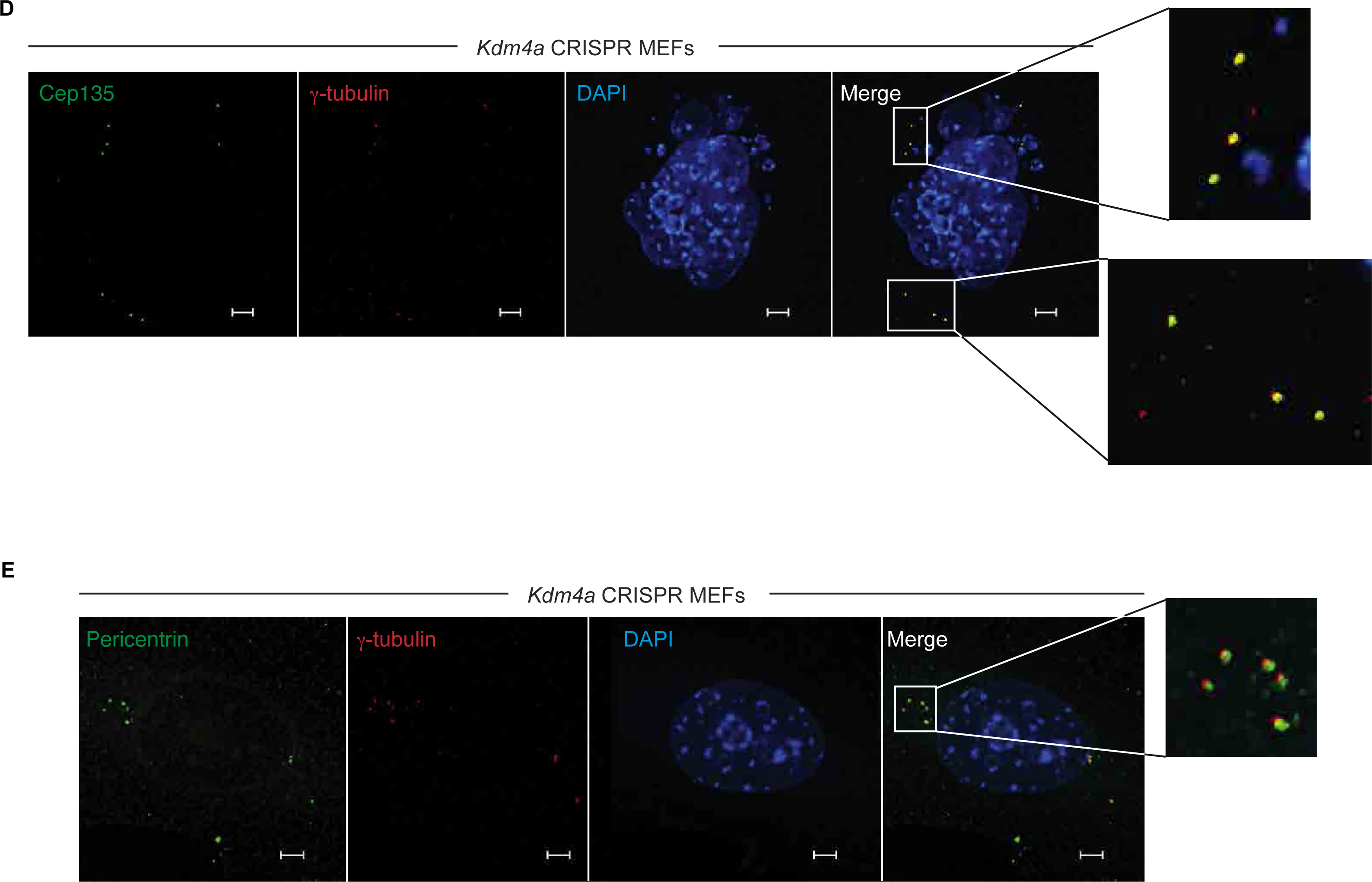
Loss of KDM4A results in supernumerary centrosomes. A Immunoblots from biochemically fractionated (nuclear and cytoplasmic fractions) *Kdm4a*-proficient and deficient MEFs probed for the indicated antibodies. B Representative images of *Kdm4a*-proficient (non-targeted) and *Kdm4a*-deficient (*Kdm4a* CRISPR) MEFs labelled with γ-tubulin (centrosome marker, red) and α-tubulin (microtubule marker, green). Nuclei are counterstained with DAPI (blue). Magnified panels show enlarged areas (white boxes) of the image for clarity. Scale bar - 10 μm. C Quantitation showing number of cells with more than 2 γ-tubulin puncta (y-axis) in *Kdm4a*-proficient (black bar) and deficient (gray bar) cells (x-axis). Each point represents values from individual experiments (n=3), each evaluating more than 500 cells. Data is represented as standard deviation (S.D.) of the mean, * p-value as indicated. D, E Representative images of *Kdm4a*-null MEFs labelled with Cep135 (centriole marker, green) (D) or pericentrin (centrosome marker, green) (E) and γ-tubulin (centrosome marker, red). Nuclei are counterstained with DAPI (blue). Scale bar – 5 μm.

### Loss of KDM4A results in pseudo-bipolar spindle formation during mitosis

Most cells that acquire extra centrosomes must ‘cope’ with these numerical anomalies, which can be especially problematic during mitosis when formation of a bipolar spindle is critical to ensure accurate chromosome segregation^14^. Given that *KDM4A* loss (*Kdm4a*-null MEFs) gave rise to supernumerary centrosomes, we next evaluated these cells through mitosis. As shown in the representative images in **Fig. 4A** and quantified in **Fig. 4B**, loss of *Kdm4a* resulted in a significant decrease in the number of bipolar mitoses with only a single centrosome at each pole. We observed a significant increase in pseudo-bipolar spindles, defined as extra centrosomes clustering at either pole **(Fig. 4A, rows 2-4)**. Formation of pseudo-bipolar spindles is a frequent coping strategy in cancer cells to enable efficient chromosome segregation when supernumerary centrosomes are present^4^. The frequency of pseudo-bipolar spindles doubled in cells with *Kdm4a* loss compared to *Kdm4a*-proficient control cells **(Fig. 4B)**. This contrasts with the percentage of monopolar **(Fig. 4B)** and multipolar spindles **(Fig. 4A, row 5 and Fig. 4B)** that showed no significant differences between *Kdm4a*-deficient and proficient cells. Again, the clustered mitotic γ-tubulin-positive foci in the pseudo-bipolar spindles were Cep135-positive confirming the presence of centrioles in these foci **(Fig. S5A)**.

**Figure 4.**
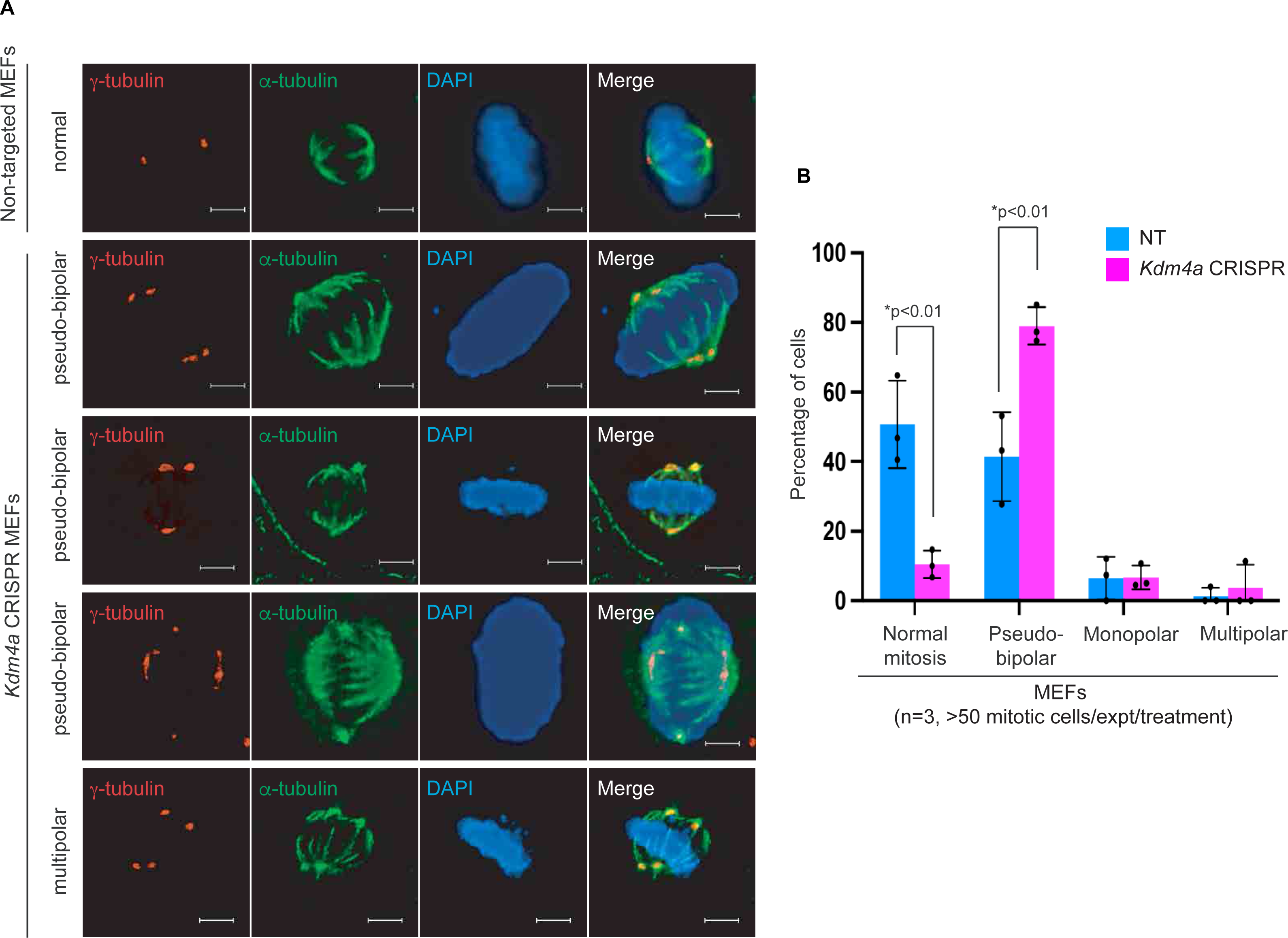
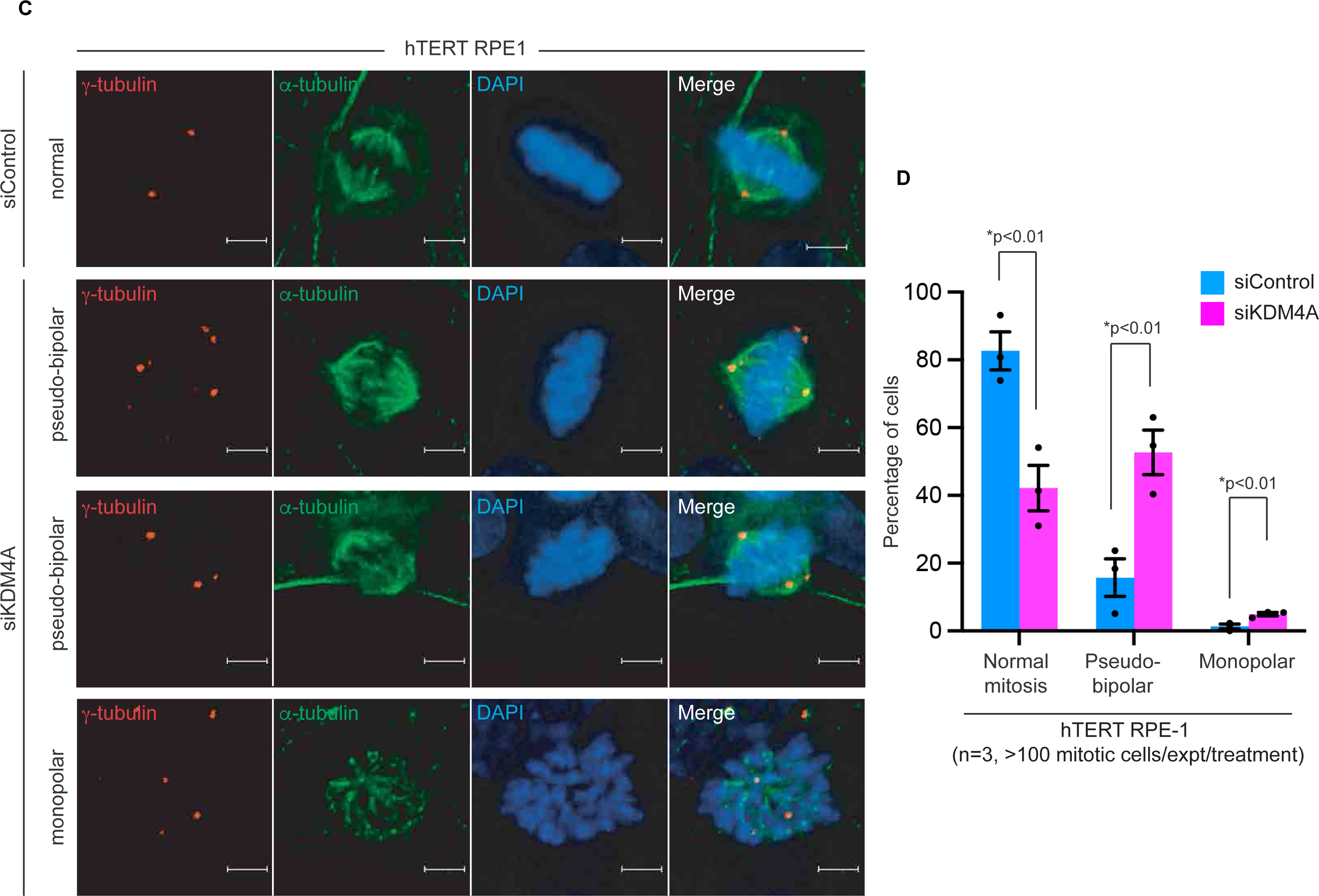
Loss of KDM4A results in aberrant mitosis. A Panel of representative mitotic images from *Kdm4a*-proficient (non-targeted) and deficient (*Kdm4a* CRISPR) MEFs labelled with γ-tubulin (centrosome marker, red) and α-tubulin (microtubule marker, green) showing normal (row 1) and aberrant (pseudo-bipolar, rows 2-4 and multipolar, row 5) spindles. Nuclei were counterstained with DAPI (blue). Scale bar – 5 μm. B Quantitation showing the percentage of mitotic defects (y-axis) in *Kdm4a*-proficient (blue bars) and deficient (pink bars) cells depicting normal, pseudo-bipolar, monopolar, and multipolar spindles (x-axis). Each point on the bar represents values from individual experiments (n=3), each evaluating more than 50 mitotic cells. Data is represented as standard deviation (S.D.) of the mean with a * p-value as indicated. C Panel of representative mitotic images of *KDM4A*-proficient (siControl) and deficient (siKDM4A) hTERT RPE-1 cells labelled with γ-tubulin (centrosome marker, red) and α-tubulin (microtubule marker, green) showing normal (row 1) and aberrant (pseudo-bipolar, rows 2-3 and monopolar, row 4) spindles. Nuclei were counterstained with DAPI (blue). Scale bar – 5 μm. D Quantitation showing the percentage of mitotic defects (y-axis) in *KDM4A*-proficient (blue bars) and deficient (pink bars) cells depicting normal, pseudo-bipolar, and monopolar spindles (x-axis). Each point on the bar represents values from individual experiments (n=3), each evaluating more than 100 mitotic cells. Data is represented as standard deviation (S.D.) of the mean with a * p-value as indicated.

To further evaluate if similar mitotic defects could be recapitulated in cells with an acute loss of *KDM4A* (compared to chronic loss using CRISPR knock out) we used an siRNA approach to knock-down *KDM4A* in hTERT RPE-1 cells **(Fig. S5B)**. Analogous to our observations in cells with chronic loss of *KDM4A* **(Fig. 4A)**, normal mitoses decreased, and pseudo-bipolar and monopolar spindles were acutely increased **(Fig. 4C, rows 2-4)**, as quantified in **Fig. 4D**. In this case, we observed a three-fold increase in pseudo-bipolar spindles accompanied by a small but significant increase in monopolar mitoses **(Fig. 4D)**. Notably, although the percentage of baseline ‘normal mitoses’ in *KDM4A*-deficient cells was different for MEFs (50%) compared to RPE-1 cells (80%), the decrease in normal mitoses with a concomitant increase in pseudo-bipolar spindles was commensurate in the genetically ‘stable’ RPE-1 cells. Together, these data suggest that cells compensate for centrosome amplification caused by loss of *KDM4A* using a clustering mechanism that enables formation of pseudo-bipolar spindles to complete mitosis in *KDM4A*-null cells.

### KDM4A contributes to genomic stability by preserving centrosome homeostasis

Evidence suggests that clustering of centrosomes as an adaptation to complete mitosis in the setting of supernumerary centrosomes still results in chromosome segregation errors and genomic instability^15^. To evaluate if chronic loss of *KDM4A* also contributes to genomic instability, we quantified the frequency of chromatin bridges and micronuclei in *Kdm4a*-null MEFs. We observed a 30-fold increase in cells with chromatin bridges in *Kdm4a*-deficient cells compared to proficient controls **(Figs. 5A-B)**. This increase was accompanied by a concordant increase in micronuclei in *Kdm4a*-null cells harboring supernumerary centrosomes, a caustic chromosome environment and a surrogate for genomic instability^4, 16^ **(Fig. 5C)**. Quantitation of micronuclei revealed more than a doubling of the percentage of cells positive for micronuclei when *Kdm4a* was lost, compared to *Kdm4a*-proficient controls **(Fig. 5D)**.

**Figure 5.**
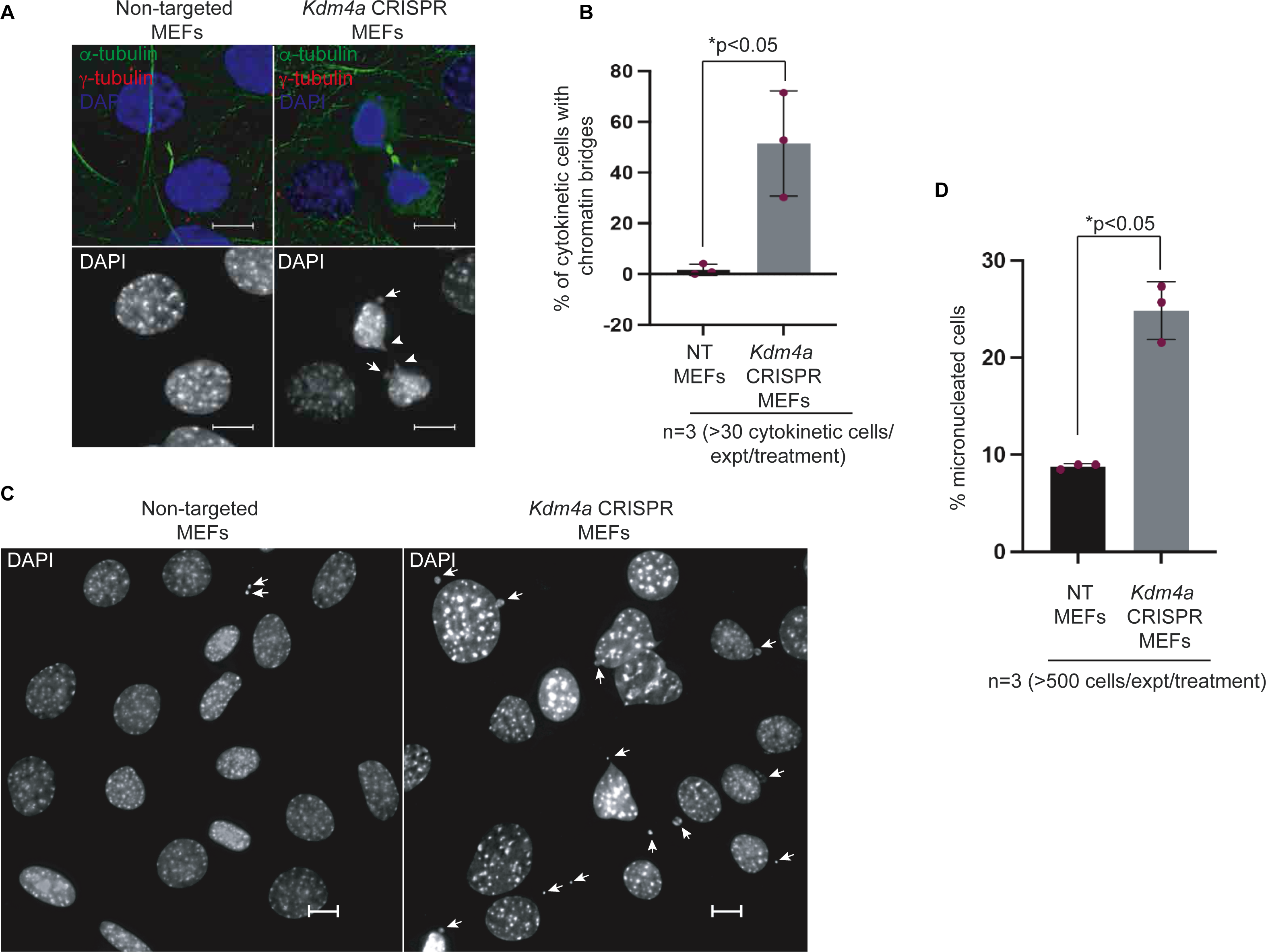
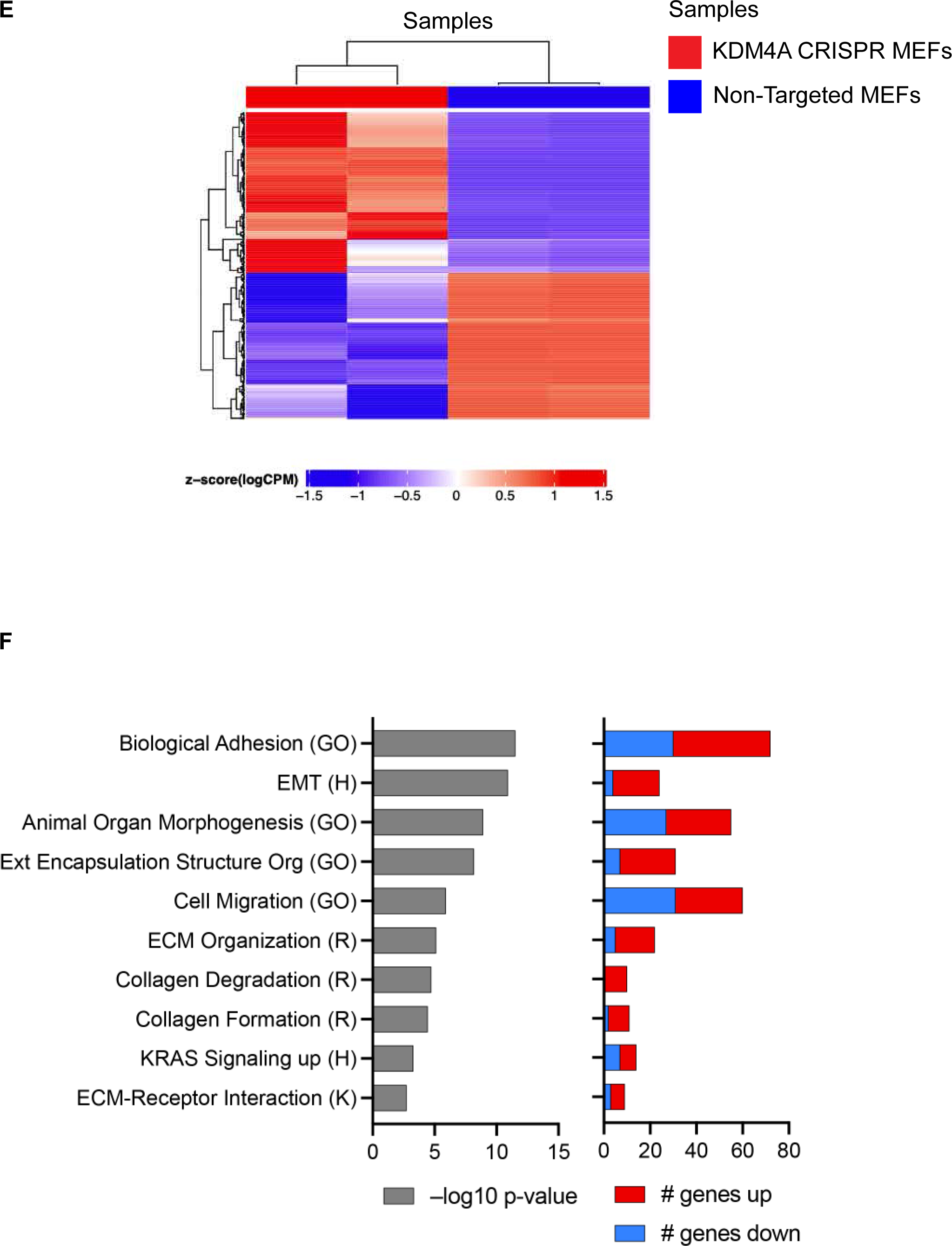
KDM4A chromatin-independent activity maintains mitotic fidelity. A Representative image showing cytokinesis in *Kdm4a*-proficient (non-targeted) and deficient (*Kdm4a*-CRISPR) MEFs. Cells were stained for α-tubulin (green), γ-tubulin (red), and nuclei were counterstained for DAPI (blue). Monochromatic grayscale images of the same are depicted to clearly show the presence of chromatin bridges. Scale bar – 5 μm. B Quantitation showing the percentage of chromatin bridges in *Kdm4a*-proficient and deficient MEFs from 3 individual experiments (individual values indicated as points on each bar graph), each evaluating more than 30 cytokinetic cells. Data is represented as standard deviation (S.D.) of the mean with a * p-value as indicated. C Representative monochromatic grayscale image showing the presence of micronuclei in *Kdm4a*-proficient (non-targeted) and deficient (*Kdm4a* CRISPR) MEFs. Scale bar – 5 μm. D Quantitation showing the percentage of micronucleated cells in *Kdm4a*-proficient and deficient MEFs from 3 individual experiments (individual values indicated as points on each bar graph) from more than 500 cells per experiment. Data is represented as standard deviation (S.D.) of the mean with a * p-value as indicated. E Heatmap of differentially expressed genes, representing a fold change exceeding 2X and FDR<0.05. F Over-representation analysis (ORA) identified enriched pathways from four MSigDB geneset collections, with the top 10 by significance shown. The number of DEGs contributing to enrichment are indicated on the right, by direction of expression.

We also asked if altered gene expression in *Kdm4a*-deficient cells could be linked to the mitotic and centrosome defects observed in these cells. RNA-seq comparing *Kdm4a*-proficient and deficient cells identified only 382 differentially expressed genes (DEGs) with 200 genes upregulated and 182 genes down regulated in *Kdm4a*-null MEFs compared to non-targeted control MEFs. Over-representation analysis (GSEA) showed that pathways enriched for these DEGs were unrelated to mitosis, centrosome biogenesis or structure **(Figs. 5E-F)**, suggesting the function of KDM4A in preserving centrosome homeostasis and genomic stability is independent of changes in gene expression caused by loss of this demethylase. The complete list of DEGs is provided in **Supplemental Table 1**.

### Enzymatic activity of KDM4A is required for mitotic fidelity

To assess if KDM4A enzymatic activity was required for maintenance of mitotic fidelity and centrosome homeostasis, we used the KDM4A pharmacologic inhibitor, JIB-04. JIB-04 is a pan-selective pharmacologic inhibitor of the Jumonji-family of demethylases that binds KDM4A and prevents its enzymatic activation without compromising its scaffolding functions^9, 17^. To specifically investigate KDM4A activity at the centrosome, we adopted a short treatment regimen of 10 μM JIB-04 for 1h, which does not increase the H3K36me3 methyl mark on chromatin as evidenced by immunoblotting **(Fig. 6A)**. While levels of H3K36me3 were unaffected in MEFs, immunofluorescence imaging of spindles in JIB-04 treated cells showed a significant increase in pseudo-bipolar spindles driven largely by centrosome splitting/fragmentation as shown in **Fig. 6B** compared to vehicle treated controls. As quantitated in **Fig. 6C**, in the presence of KDM4A inhibitor we saw a 6-fold increase in pseudo-bipolar spindles relative to vehicle controls.

**Figure 6.**
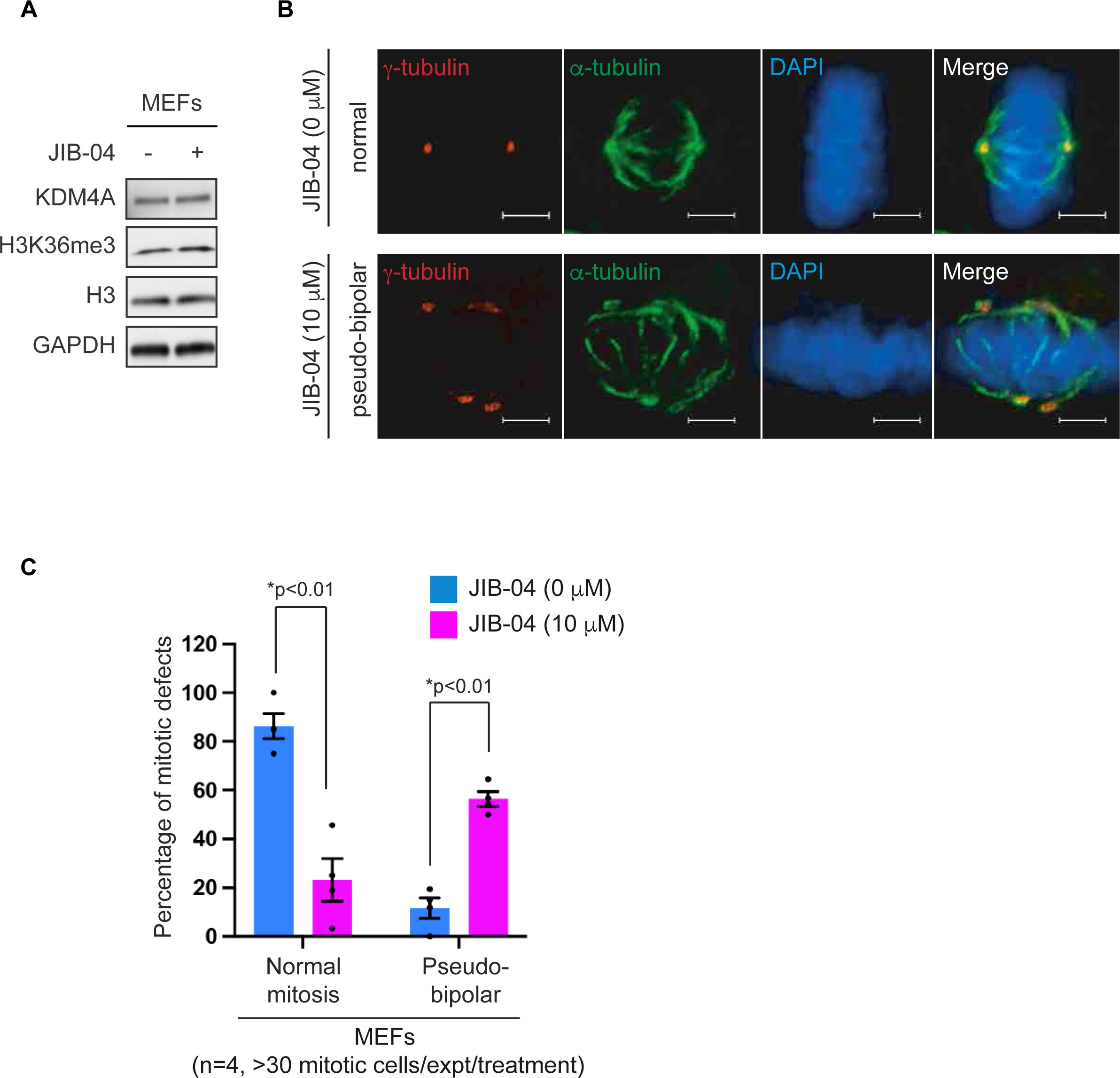
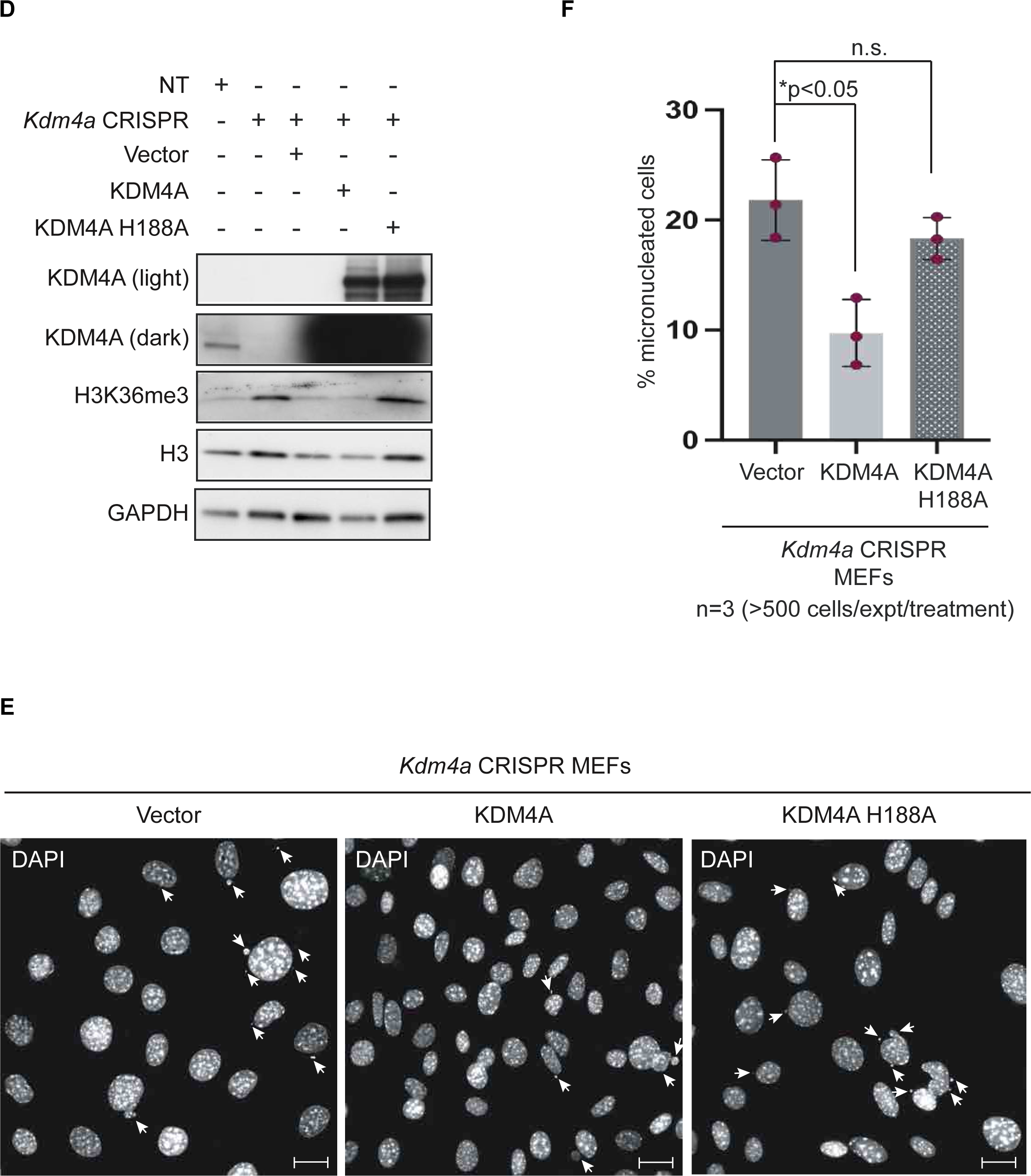
Loss of KDM4A enzymatic activity results in genomic instability. A Immunoblots of lysates generated from *Kdm4a*-proficient MEFs treated with KDM4A-inhibitor JIB-04 (10 μM, 1h) probed for the indicated antibodies. B Representative images of mitosis from *Kdm4a*-proficient MEFs treated with vehicle (row 1) or 10 μM JIB-04 (KDM4A inhibitor, rows 2-3) labelled for γ-tubulin (centrosome marker, red) and α-tubulin (microtubule marker, green) showing aberrant (pseudo-bipolar) spindles. Nuclei were counterstained with DAPI (blue). Scale bar – 5 μm. C Quantitation showing the percentage of mitotic defects (y-axis) in vehicle treated (0 μM, blue bars) and 10 μM JIB-04 treated (pink bars) MEFs depicting normal and pseudo-bipolar spindles (x-axis). Each point on the bar represents values from individual experiments (n=4), each evaluating more than 30 mitotic cells. Data is represented as standard deviation (S.E.M.) of the mean with a * p-value as indicated. D Immunoblots from *Kdm4a*-null MEFs rescued with wild-type KDM4A and catalytically dead H188A KDM4A probed for the indicated antibodies. E Representative monochromatic grayscale images showing the presence of micronuclei in *Kdm4a*-null MEFs rescued with wildtype KDM4A and catalytically dead H188A KDM4A. Scale bar – 5 μm. F Quantitation showing the percentage of cells with micronuclei in *Kdm4a*-deficient MEFs rescued with wild-type KDM4A or catalytically dead H188A KDM4A from 3 individual experiments (individual values indicated as points on each bar graph) from more than 500 cells per experiment. Data is represented as standard deviation (S.D.) of the mean with a * p-value as indicated.

These data showing enzymatic activity of KDM4A regulating mitotic fidelity prompted us to investigate if micronuclei formation could be rescued in *Kdm4a*-null cells re-expressing an enzymatically proficient (wild-type KDM4A) or catalytically deficient (H188A) mutant KDM4A. Immunoblots confirmed expression of KDM4A in the rescue with concordant decrease of H3K36me3 exclusively in cells expressing KDM4A wild type (enzymatically proficient) but not the catalytically dead H188A KDM4A (enzymatically deficient) **(Fig. 6D)**. Importantly, cells rescued with wild type KDM4A demonstrated decreased micronuclei formation. However, cells expressing catalytically dead KDM4A failed to rescue the micronuclei phenotype exhibiting rates comparable to the *Kdm4a*-null cells expressing empty vector (control) **(Fig. 6E-F)**. These data identify a role for the demethylase activity of KDM4A in preserving genomic stability.

## DISCUSSION

In this study, we have identified a spatiotemporal role for KDM4A at the centrosomes during mitosis, where it is required for mitotic fidelity and genomic stability. We showed KDM4A localized at MTOCs in dividing cells and at centrosomes in interphase cells. Loss of *KDM4A* caused supernumerary centrosomes, with the surplus centrosomes clustered at pseudo-bipolar spindles in cells progressing through mitosis. Maintenance of centrosome homeostasis by KDM4A required this enzyme’s catalytic activity: loss or inhibition of its enzymatic activity decreased mitotic fidelity, resulting in pseudo-bipolar spindles, chromosome segregation defects and genomic instability. In summary, our data revealed a new role for this demethylase at the centrosome, adding KDM4A to the growing list of dual-functioning chromatin modifying enzymes required for mitotic spindle integrity and genomic stability.

KDM4A demethylates histones by removing methyl marks from lysine 36 (H3K36me2/3) and lysine 9 (H3K9me2/3) on histone tails^9^. Although primarily a nuclear protein, KDM4A was identified in the cytoplasmic fractions of cells, where it interacted with the translation initiation complex to regulate protein synthesis^18^. Our data reveals an additional, non-chromatin role for KDM4A as a regulator of centrosome integrity and mitotic fidelity. Direct immunofluorescence analysis and single-molecule super-resolution imaging of KDM4A showed focally localized signal at centrosomes during cell division, with minimal KDM4A staining on chromatin during mitosis, corroborating the previous report of low expression levels observed in G2/M^19^.

Our single-molecule super-resolution data further revealed a spatial distribution of KDM4A that appeared to uniformly encompass the centrosome structure, lacking specific centriolar localization (on either mother or daughter), suggesting that perhaps KDM4A is a component of the pericentriolar material around the centrioles. While chromatin modifying enzymes have not previously been identified in mass spectrometry-based proteomic screens characterizing centrosome proteins^20, 21, 22^ this dichotomy likely arises from use of analytical pipelines that exclude ‘contaminating’ fragments, or use more targeted proteomic approaches, all of which could result in an exclusion of ‘nuclear’ chromatin proteins. Our data establishing KDM4A localization and function at the centrosome now pave the way for future studies aimed at understanding the contributions of non-canonical centrosome proteins, such as chromatin modifying enzymes, in the maintenance and activity of this complex structure.

We showed acquisition of supernumerary centrosomes was a consequence of chronic loss of *KDM4A* (CRISPR-induced). Extra centrosomes pose a unique challenge to bipolar spindle formation during mitosis, and cells must develop strategies for managing this excess for mitotic success^23^, as formation of multipolar spindles triggers elimination of these cells from the population^23^. Indeed, multipolar spindles were seen at exceedingly low frequencies in the setting of chronic *KDM4A* loss, likely due to the ability of surviving cells to cluster extra centrosomes at either pole during mitosis to compensate for the presence of supernumerary centrosomes. Multiple mechanisms exist for the generation of amplified centrosomes in cells ranging from overamplification, centrosome fragmentation, or failure to complete cytokinesis^14^. Although the supernumerary centrosomes observed in cells with a chronic loss of *Kdm4a* could arise via any of these approaches, our data with the KDM4A inhibitor, showing increased γ-tubulin puncta at the spindle within an hour of treatment with the inhibitor (JIB-04), is suggestive of a mechanism independent of aberrant S-phase centrosome replication. Thus, in cells treated with JIB-04 (KDM4A inhibitor) centrosome/centriolar fragmentation or even a loss of cohesion between the centrioles could result in delayed progression through mitosis, triggering cell death. In contrast, the generation of amplified centrosomes in cells with a chronic loss of *KDM4A* could in part be driven by replication defects during S-phase that could result in uncoupling of centrosome and cell cycle. A systematic evaluation of each process would be required to identify conclusively a mechanism for the amplified centrosomes in the absence of this demethylase.

Loss of *KDM4A* resulted in altered gene expression, as expected. However, importantly the differentially expressed genes enriched for pathways unrelated to spindle formation, cell division or centrosome biogenesis/ homeostasis, consistent with KDM4A regulation of centrosome homeostasis occurring via mechanism(s) other than reprogramming of gene expression. However, one caveat to this interpretation is that active transcription is dramatically reduced during mitosis with the bulk of gene expression changes occurring during interphase^24^. Given that the bulk RNA-seq analysis were not specific to a mitotic population, the differential expression changes observed could be attributed to the non-mitotic populations. Nonetheless, these data corroborate previous studies that established a role for chromatin modifiers on microtubules of the mitotic spindle i.e., SETD2 in methylating microtubules^25^ and PBRM1 in reading this SETD2 methyl mark^26^, which similarly showed few gene changes and did not enrich for genes in pathways related to mitosis. In addition, the rapid time frame within which multiple puncta/clustered centrosomes became apparent in KDM4A inhibitor (JIB-04) treated cells, irrespective of how they develop, further hints to a likely chromatin-independent role for KDM4A in driving centrosome abnormalities.

An increased incidence of chromosome segregation defects, including chromatin bridges, and an exacerbation of micronuclei formation in *KDM4A*-null cells containing supernumerary centrosomes supports a role for KDM4A in maintaining genomic stability. The increase in centrosome numbers and ensuing clustering of these during mitosis is often associated with errors in microtubule attachments to the kinetochore, which could be a significant contributor to chromosome segregation defects^7^. Our data establishing a functional role for KDM4A in centrosome homeostasis presents a novel pathway to genomic instability via dysregulated centrosome biogenesis. Further studies will be necessary to understand the mechanistic underpinnings and molecular targets of KDM4A enzymatic activity at the centrosome.

## METHODS

### Cell Culture

All cell lines used in the study were cultured at 37°C and 5% CO_2_. Immortalized human retinal pigmented epithelial (hTERT-RPE1) cells (gift from Dr. Gregory Pazour, University of Massachusetts Medical School, Worcester, MA) were cultured in DMEM/F-12 (ThermoFisher Scientific, Cat. #11320-082). Human kidney cells (HKC) and human embryonic kidney cells (HEK293T) were cultured in Dulbecco’s Modified Eagle Medium (DMEM, ThermoFisher Scientific, Cat. # 11965126). Mouse embryonic fibroblasts (MEF) were cultured in DMEM phenol red free media (ThermoFisher Scientific, Cat. # 21063-029) supplemented with sodium pyruvate (Gibco, Cat. # 11360070), GlutaMAX^TM^ supplement (Gibco, Cat. # 35050061) and Blasticidin S (ThermoFisher Scientific, Cat. # A1113903). All cell lines were supplemented with 10% Fetal Bovine Serum (FBS, Sigma-Aldrich, # F2442). Cell lines were routinely monitored for mycoplasma and were confirmed negative before use for experiments.

### Transfections and Drug Treatments

#### siRNAs

ON-TARGETplus SMARTpool siControl (Cat. # D-001810-01-20) and siKDM4A (Cat. # M-004292-01) were obtained as lyophilized powder from Dharmacon (Revvity). siRNA was resuspended in 250 µL of 1X siRNA buffer (Dharmacon, Cat. #B-002000-UB-100) to obtain a 20 µM final stock. Cells were transfected with siC, siKDM4A at a final concentration of 10 nM using DharmaFECT1 transfection reagent (Dharmacon, Cat. # T-2001-03) for 48h.

#### KDM4A constructs

For overexpression studies, the HA-KDM4A (Addgene, Plasmid # 24180) construct was transfected using Lipofectamine2000 (ThermoFisher Scientific, Cat # 52932) according to the manufacturer’s protocol. Rescue cell lines in *Kdm4a*-null MEFs with wildtype and catalytically dead KDM4A were generated with constructs Lentizeo-RFP, Lentizeo-KDM4A and Lentizeo-KDM4A-H188A (kind gifts of Dr. Fred van Leeuwen, University of Amsterdam, Netherlands).

#### JIB-04 treatment

JIB-04 was purchased from Selleckchem (Cat. # S7281) and was resuspended in DMSO to obtain 100 mM stocks. Cells were plated and treated at 70-80% confluency with JIB-04 at 10 μM for 1h. Cells were subsequently harvested to prepare whole cell extracts for immunoblotting or fixed with methanol for immunofluorescence.

### Generation of Stable Cell Lines

#### GFP-centrin RPE-1 cells

To generate hTERT-RPE1 stably expressing GFP-centrin, a centrosome marker, cells were transfected with pEGFP-C1-Centrin-1 plasmid (Addgene, Plasmid # 72641) using Lipofectamine2000 according to the manufacture’s recommendations. Cells expressing the construct were selected using G418 (Geneticin, ThermoFisher Scientific Cat. # 10131027) at 800 µg/ml, and single colonies picked and validated by immunoblotting and immunofluorescence.

#### Kdm4a CRISPR MEFs

*Kdm4a* was knocked out in MEFs using CRISPR-Cas9 gene editing. sgRNA guides (sgKdm4a) co-expressing Cas9 were purchased from GenScript. Three sgRNA constructs were co-transfected into MEFs and selected using puromycin (2 μg/ml) (ThermoFisher Scientific, Cat. # A11138-03). Single colonies were selected and validated by immunoblotting to confirm *Kdm4a* knockout. The sequences for the guides used are as follows: sgKdm4a#1-TAGATCATCAATATCGTCGT, sgKdm4a#2 – GATCTTGCGGAACTCACGAA, sgKdm4a#3 – CGGCCGGCTGAAGACCATCC.

#### Kdm4a CRISPR rescue cell lines

The MEFs with *Kdm4a* knockout were rescued using the Lentizeo-RFP, Lentizeo-KDM4A and Lentizeo-KDM4A-H188A constructs. Lentivirus was generated in HEK293FT cell (ThermoFisher Scientific, # R70007) co-transfected with helper constructs, PAX2 plasmid (Addgene, Plasmid # 35002) and pMD2.G plasmid (Addgene, Plasmid # 12259). The conditioned media was collected and filtered 48h post transfection and used to infect the *Kdm4a*-null MEFs using polybrene (Millipore Sigma, Cat. # T1003-G) at 10 µg/ml. Cells were expanded and selected using Zeocin (ThermoFisher Scientific, Cat. # R25001) at 800 µg/ml. Single colonies were selected and screened by immunoblotting.

### Antibodies for Immunofluorescence and Immunoblotting

Antibodies were procured from the following sources: anti-KDM4A (1:1000, Sigma Aldrich, Cat. # HPA007610), anti-γ-tubulin (1:4000, Sigma Aldrich, Cat. # T5326), anti-α-tubulin (1:4000, Sigma Aldrich, Cat. # T6199), anti-α-tubulin (1:1000, ThermoFisher Scientific, Cat. # PA5-19489), anti-α-tubulin DM1A (1:5000, DSHB, Cat. # T6199), anti-α-tubulin (1:1000, Abcam, Cat. # ab6161), anti-HA (1:1000, Cell Signaling, Cat. # 3724S), anti-lactate dehydrogenase (LDH) (1:2000, Abcam, Cat. # ab47010), anti-lamin A/C (1:2000, Cell Signaling, Cat. # 2032S), anti-SETD2 (1:1000, Sigma Aldrich Cat. # HPA-042451-100), anti-H3K36me3 (1:1000, Active Motif, Cat. # 61101), anti-H3 (1:5000, Cell Signaling, Cat. # 4499), anti-CEP135 (1:200, Abcam, Cat. # ab75005), anti-pericentrin (1:4000, Abcam, Cat. # ab4448), anti-CP110 (1:500, Millipore Sigma, Cat. # MABT1354), anti-centrobin (1:200, Abcam, Cat. # ab70448), anti-centrin 2 (1:200, Millipore Sigma, Cat. # 04-1624), anti-GAPDH (1:5000, Proteintech, Cat. # 10494-1-AP). HRP conjugated secondary goat anti-mouse (Cat. # 170-6516), and goat anti-rabbit (Cat. # 170-6515) antibodies were used at 1:2000 for immunoblotting and purchased from Biorad.

### Immunoblotting

To make whole cell lysates, cells (70–80% confluent unless otherwise noted) were washed and collected in ice-cold PBS and pelleted by centrifugation at 4°C. Pellets were either flash frozen in liquid nitrogen and stored at −80°C for future use or resuspended in 1X lysis buffer (25 mM Tris, pH 8.0, 300 mM NaCl, 1 mM EDTA, 1% NP40). 1X complete protease inhibitor cocktail (Roche, Cat. # 04693132001), 1 mM sodium orthovanadate (Na_3_VO_4_, Sigma Aldrich, Cat. # S6508), and 1 mM PMSF (Sigma Aldrich, Cat. # P7626) were added fresh to the lysis buffer prior to use. The cellular extracts were sonicated for 10 cycles (30 seconds on and 30 seconds off per cycle) using a Diagenode Bioruptor 300 and the extracts centrifuged at maximum speed for 10 min at 4°C. The pellet was discarded, and supernatant collected as whole cell extract (WCE). Whole cell extracts were subjected to BCA-protein assay to quantify and normalize protein levels. Soluble proteins were subject to immunoblot analysis using 4-15 % SDS-PAGE gels (Bio-Rad, Cat. # 64557025) followed by transfer to PVDF membrane. Membranes were blocked for 1h in 5% non-fat milk and incubated with primary antibodies overnight at 4°C. Membranes were subsequently washed 3 times with TBST, and primary antibodies were tagged with horseradish peroxidase (HRP) conjugated goat anti-mouse and goat anti-rabbit secondary antibodies by incubation for 1h at RT. Membranes were then washed 3 times with TBST (10mins each) and visualization performed using ECL and ECL Prime (ThermoFisher Scientific, Cat. # 32106 and Amersham, Cat. # RPN2232, respectively).

### Sub-Cellular Fractionation

For subcellular fractionation, flash frozen cell pellets were resuspended in “Solution A” (sucrose- 5.47g, 10mM Tris-HCl, pH 8.0, 3.0mM CaCl_2_, 0.1mM EDTA, 2mM magnesium acetate, 0.5% NP40). The solution was well-mixed by pipetting several times and incubated on ice for 10 min to ensure complete lysis. After centrifugation at 4°C at 1500 g for 5 min, the supernatant was collected as crude cytoplasmic fraction and the pellet as crude nuclei. The crude cytoplasmic fraction was recentrifuged at full speed for 10 min to pellet any suspended impurities and the supernatant collected as ‘cytoplasmic fraction’. The crude nuclei were washed with “Solution B” (sucrose 5.47g, 10 mM Tris-HCl, pH 8.0, 3.0 mM CaCl_2_, 0.1 mM EDTA, 2 mM magnesium acetate) and the pellet obtained from centrifugation at 4°C at 1500 g for 5 min was lysed in 1X lysis buffer (25 mM Tris, pH 8.0, 300 mM NaCl, 1 mM EDTA, 1% NP40) and sonicated for 10 cycles (30 seconds on and 30 seconds off) using a Diagenode Bioruptor 300 to ensure complete nuclear lysis. After centrifugation at 13,000 rpm at 4°C for 10 min, the pellet was discarded, and supernatant collected as ‘nuclear fraction’. All lysis and wash buffers contained 1X complete protease inhibitor cocktail, 1 mM sodium orthovanadate and 1 mM PMSF. Nuclear and cytoplasmic fractions were subjected to BCA protein assay to quantify and normalize protein levels and then subjected to SDS-PAGE and immunoblot analysis.

### Immunoprecipitation

ChromoTek GFP-Trap Magnetic Agarose beads (Proteintech, Cat. # gtma20), and Pierce Magnetic A/G beads (ThermoFisher Scientific, Cat. # 88803) used for immunoprecipitation assays were purchased from the sources detailed. Whole cell extracts (800-1000 µg) were first pre-cleared with Magnetic A/G beads for 3h with constant rocking. Beads were discarded and the pre-cleared lysates were incubated with necessary antibodies overnight at 4°C with constant rocking. Magnetic beads were then added to these samples and further incubated for 1h at 4°C with constant rocking. The immunoprecipitated samples were washed 5 times (10 mins each) with 1X lysis buffer (25 mM Tris, pH 8.0, 300 mM NaCl, 1 mM EDTA, 1% NP40, 1X complete protease inhibitor cocktail, 1 mM sodium orthovanadate and 1 mM PMSF). Beads were then resuspended in 2X denaturing sample loading buffer, heated for 10 min at 95°C and loaded on an SDS-PAGE gel for immunoblotting analysis. In the case of GFP-TRAP experiments, pre-clearing was not necessary.

### Direct Immunofluorescence

Cells for immunofluorescence analysis of centrosomes were fixed using pre-chilled (−20°C) methanol (ThermoFisher Scientific Cat. # A412P-4) for 10 minutes at −20°C. Alternately, cells used for genomic stability analysis were fixed using 4% paraformaldehyde (PFA) (Electron Microscopy Sciences, Cat. # 15714) for 15 min at RT. This was followed by permeabilization using 0.5% Triton-X for 10 min. 3.75% bovine serum albumin (BSA) (Sigma Aldrich, Cat. # A8412) was used as the blocking buffer and to dilute both primary and secondary antibodies. Samples were incubated in primary antibodies overnight with gentle rocking, washed 3 times (10 min each) followed by incubation with secondary antibodies for 1h at room temperature in 3.75% BSA. Alexa Fluor 488, 546, 647 anti-mouse (Life Technologies, Cat. # A11001, A11003 and A21235 respectively) and 488, 546, 647 anti-rabbit (Life Technologies, Cat. # A11034, A11010 and A21245, respectively) were used as secondary antibodies. Cells were washed 3 times (10 min) and post fixed with 4% PFA followed by counterstaining of nuclei using DAPI (1:4000 for 10 min, ThermoFisher Scientific, Cat # 62248). Coverslips were mounted onto glass slides and direct fluorescence of mitotic cells and genomic instability visualized using a deconvolution microscope (ECLIPSE Ti2 Inverted Microscope, NIKON) at 60X magnification.

### Single Molecule Super Resolution Immunofluorescence

#### PDMS (Polydimethylsiloxane) Microfluidic Chip Preparation

Prior to experiments, PDMS chips with microfluidic channels were prepared using SU8 molds^27, 28, 29, 30, 31^. The SU8 molds were prepared by spin-coating approximately 5 mL of the photoresist (Kayaku Advanced Materials, Inc., Cat. # Y131273) onto a silicon wafer (University Wafer, Inc., Cat. # 444) by first ramping to 500 revolutions per min (rpm) at 100 rpm/s acceleration, then holding at 500 rpm for 10s, then ramping to 3000 rpm at an acceleration of 300 rpm/s and holding at this speed for 30 s. These molds were then placed in an oven at 65°C for 10 min before being placed on a hot plate at 95°C for 30 min. Film photomasks (Micro Lithography Services Ltd.) with opening dimensions of 1 cm x 1-3 mm were placed on the silicon wafers and exposed to 18 W of UV light under vacuum (Vacuum UV-Exposure Box, Gie-Tec GmbH, Cat. # 140010) for 27 s. The wafers were then placed in the 65°C oven for 1 min before being placed on the 95°C hot plate for 10 min. The wafers were then gently swirled in a beaker of approximately 20 mL SU8 Developer (mr-Dev 600, Kayaku Advanced Materials, Inc.) for 15 min to dissolve photoresist in the areas that had not been illuminated. This procedure generated molds that had 100 µm high ridges. PDMS (SYLGARD 184 Silicone Elastomer Kit, Dow Inc.) was then added to these SU8 molds to generate PDMS chips with microfluidic channels 1 cm long, 1-3 mm wide, and 100 µm high. Approximately 20-25 mL of the combined PDMS product was poured onto the molds placed in a petri dish (Fisher Scientific, Cat. # FB0875713A) with pins positioned at either end of the channel to create holes for tubing. The chips were left to cure for at least 48h, then removed from the petri dish, and cut into approximately 1 cm x 2 cm chips. The chips were stored at RT protected from dust until used.

#### Immunolabeling

Cells were fixed with 100% methanol on plasma-cleaned coverslips (#1.5H, 22 x 22 mm, 170 ± 5 µm, Thorlabs, Cat. # CG15CH). The slides were then submerged and stored in PBS at 4**°**C for up to 6 months in 6 well plates (Falcon™ Polystyrene Microplates, Cat. # 08-772-1G). Prior to permeabilization, a PDMS chip was attached to the top of the coverslip with General Purpose 5- Minute Epoxy (Thorlabs, Cat. # G14250) to create a small microfluidic chamber with holes on each end of the channel. Cells were permeabilized for 10 min using 0.05% (v/v) Triton-X 100 in PBS. They were then washed 2x with PBS and blocked using 3% bovine serum albumin (BSA, A2058, Sigma-Aldrich) in PBS for 1h at RT. The cells were incubated with rabbit anti-KDM4A (1:500), mouse anti-centrobin (1:300), and rat anti-tubulin (1:500) antibodies diluted in 3% BSA in PBS at 4°C overnight, and then washed 3 times with PBS with 10 min incubation for each wash. The cells were then washed with washing buffer (1x, Massive Photonics, Cat. # 2301), incubated with donkey anti-mouse + Docking site 1 (Massive Photonics, Cat. # 2301, 1:100), donkey anti-rabbit + Docking site 2 (Massive Photonics, Cat. # 2301, 1:100), and goat anti-rat Alexa Fluor 647 (Invitrogen, A-Cat. # 21247, 1:1000) antibodies diluted in antibody incubation buffer (Massive Photonics, Cat. # 2301) for 1h at RT shielded from light, then washed 3 times with washing buffer (1x, Massive Photonics). Fluorescent beads (Invitrogen, Cat. # F8807, 1:50,000) diluted in MilliQ water were added while the sample was on the microscope and washed out with PBS once beads adhered to the coverslips after approximately 5 min.

#### Optical Setup

The optical setup was built around a conventional inverted microscope (IX83, Olympus) **(Fig. S2)**^12, 32^. Excitation lasers (560 nm and 642 nm, both 1000 mW, MPB Communications) were circularly polarized (LPVISC050-MP2 polarizers, Thorlabs; 560 nm: Z-10-A-.250-B-556 and 642 nm: Z-10-A-.250-B-647 quarter-wave plates, both Tower Optical) and filtered (560 nm: FF01-554/23-25 excitation filter, 642 nm: FF01-631/36-25 excitation filter, both Semrock), and expanded and collimated using lens telescopes. Collimated light was introduced into the back port of the microscope and focused by a Köhler lens to allow for wide-field epi-illumination. The lasers were toggled with shutters (VS14S2T1 with VMM-D3 three-channel driver, Vincent Associates Uniblitz). The sample was positioned on an xy translation stage (Physik Instrumente, Cat. # M26821LOJ) and an xyz piezoelectric stage (Physik Instrumente, Cat. # P-545.3C8H). The emission from the sample was collected using a high numerical aperture (NA) objective (100x, NA 1.45, Olympus, Cat. # UPLXAPO100XO) and filtered (ZT405/488/561/640rpcV3 dichroic; ZET561NF notch filter; and ZET642NF notch filter, all Chroma) before entering a 4f imaging system. The first lens of the 4f imaging system (f = 80 mm, Thorlabs, Cat. # AC508-080-AB) was placed one focal length from the intermediate image plane in the emission path. A dichroic mirror (Chroma, Cat. # T660lpxr-UF3) was placed after the first 4f lens to split the light into two different spectral paths, where far red light (“red channel”) was transmitted into one optical path and greener light (“green channel”) was reflected into the other optical path. In order to reshape the PSF of the microscope to encode the axial position (z) of the individual fluorophores, transmissive dielectric double-helix (DH) phase masks (red channel: DH-12 phase mask, 670 nm with diameter of 2.484 mm, Double-Helix Optics, LLC; green channel: DH-1 phase mask, 590 nm with diameter of 2.484 mm, Double-Helix Optics, LLC) were placed one focal length after the first 4f lens in each path and another 4f lens was placed one focal length after the phase masks in both paths^33, 34, 35, 36, 37, 38,39^. Bandpass filters (red channel: ET700/75m; green channel: ET605/70m, both Chroma) were placed in the paths between the phase masks and the second 4f lenses. The second 4f lenses then focused the light onto an EM-CCD camera (iXon Ultra897, Andor) placed one focal length away from the second 4f lenses.

#### 3D Exchange-PAINT imaging

To facilitate calibration of the engineered PSFs and registration between the two channels, a solution of fiducial beads (Tetraspeck, T7280, 0.2µm, Invitrogen) were diluted 1:5 in 10% polyvinyl alcohol (Mowiol 4-88, 17951, Polysciences Inc.) in MilliQ water and spun-coat onto plasma-cleaned coverslips (#1.5H, 22 x 22 mm, 170 ± 5 µm, Thorlabs, Cat. CG15CH). For calibration of the PSFs, images were acquired while scanning axially over 2 μm and 12 μm axial ranges with 50 nm and 100 nm steps, respectively, using the piezoelectric xyz translation stage. For registration between the two channels, frames were acquired and averaged at ten different xy positions across the coverslips with at least 5 fiducial beads in each frame. Dark frames (400) were collected with the camera shutter closed before image acquisition, and the averaged intensity was subtracted from the calibration, registration, single-molecule, and fiducial bead data before further analysis.

For diffraction-limited imaging, cells were imaged using laser intensities of 0.3 W/cm^2^ for the 642 nm laser and 1.2 W/cm^2^ for the 560 nm laser. For single-molecule imaging of KDM4A and centrobin, sequential DNA-based point accumulation for imaging in nanoscale topography (DNA-PAINT) called Exchange-PAINT was used ^13^. Using a microfluidic pump, Cy3B-conjugated imager strands 1 and 2 (0.05 nM, Massive Photonics, Cat. # 2301) were flown into the microfluidic chamber sequentially with washing buffer (1x, Massive Photonics) flown in between. Frames (100,000 for each target) were acquired for each imager strand using a laser power of ∼0.7 kW/cm^2^ for the 560 nm laser. An exposure time of 0.125 s was used with a calibrated EM gain of 183 and conversion gain of 4.41 photoelectrons/ADC count.

#### Analysis of 3D single-molecule super-resolution data

Calibration and analysis of fiducial bead stacks and data and analysis of single-molecule images were performed using the open-source Easy-DHPSF software ^40^ (https://sourceforge.net/projects/easy-dhpsf/). For the fiducial bead data acquired in the “red” channel using the 12 µm DH phase mask, the Easy-DHPSF code was altered as follows: in *easy_dhpsf.m*, s.sigmaBounds = [1.2 3.0], s.lobeDistance = [38 45], and s.boxRadius = 21. In *F_calDHPSF.m*, templateSize = 70. Registration of the two channels was performed using a transformation matrix from the fiducial bead “red” channel to the single-molecule “green” channel using the MATLAB function *imregtform*. Transformed fiducial bead localizations were used to drift correct the single-molecule data in the “green” channel using a custom cubic spline fitting function for the fiducial bead data. Localizations were rendered with Vutara SRX (Bruker), where each localization was represented by a 3D Gaussian with 25 nm diameter and with variable opacities set to best visualize the localization density. The localizations were filtered to remove localizations with xy and z localization precision, σ_xy_ and σ_z_, above 20 nm and 30 nm, respectively. For quantification of the KDM4A distributions, KDM4A localizations were manually isolated and exported from Vutara SRX in .mat files. These distributions were then imported into MATLAB and the 1/e^2^ diameters of each of the five distributions from three different cells were calculated independently in the x,y, and z directions **(Fig. S4)**.

## RNA-Seq Analysis

Total RNA was extracted from KDM4A-proficient and deficient MEFs using TRIzol reagent (ThermoFisher Scientific, Cat. # 15596026) and the resulting RNA cleaned using RNeasy Mini Kit (Qiagen, Cat. # 74104) according to the manufacturer’s instructions. Quality of extracted RNA was determined using the RNA Tapestation 4200 (Agilent) and quantified using Qubit Fluorometer (ThermoFisher Scientific). The RNA libraries were prepared and sequenced at the University of Houston Sequencing and Editing Core facility per their standard protocols. The prepared libraries were pooled and sequenced using the NextSeq 500 (Illumina) platform. All samples were prepared and run in duplicate. RNA-seq paired-end reads were trimmed using cutadapt^41^ 1.18 and fastQC^42^ v0.11.9. Mapping was done with Homo_sapiens.GRCh38.101.gtf^43^ as a reference genome. Some low-quality samples were found after trimming and mapping quality control with the multiqc^44^ utility version 1.8. The samples 15743-1_s14 and 17509-1_S13 were removed from further analysis. Differential expression analysis was done with use of the edgeR^45^ package version 3.32.1 and EDAseq^46^ 2.24.0. An FDR cutoff of 0.05 was selected and fold change cutoffs of 2.0; LRT (likelihood ratio test) RUVr (remove unwanted variation) upper quartile normalization was used.

Over-representation Analysis (ORA) was performed to detect enrichment of gene sets corresponding to pathways and biological processes, based on differential gene expression. Using the Hallmark compendium (v7.5) and the Molecular Signature Database methodology (MSigDB)^47^, a hypergeometric test was used to assess the enrichment with significance achieved at adjusted p-value<0.05.

## ACKNOWLEDGEMENTS

The authors would like to thank the members of the Dere lab for their thoughtful comments and feedback. This work was supported by research funding from the Cancer Prevention Research Institute of Texas (RP220332), Department of Defense CDMRP KCRP (KC210123) to RD; NIH/NCI (R01-CA275082) to FM and RD; Department of Defense (W81XWH2110786) to FM; Cancer Prevention Research Institute of Texas (R86390), the Welch Foundation (C-2064-20210327), the NIH/NIGMS (R00GM134187) to AKG; the Rice University Provost’s TMC Collaborator Seed Fund Program (PTC2401) to AKG and RD; the Welch Foundation (I-1878) to EDM; NIH/NCI (R35CA231993) and NIH/NIEHS (P30ES030285) to CLW. SLG ad CC were partially supported by the Cancer Prevention Research Institute of Texas (CPRIT) (RP210227, RP200504), NIH/NCI P30 shared resource grant (CA125123), NIH/NIEHS center grants (P30 ES030285, P42 ES027725), NIH/NIMHD (P50MD015496), and NIH/NIMH (R01MH134392).

## AUTHOR CONTRIBUTIONS

PC performed the biochemical and imaging experiments with contributions from XW (fluorescent imaging), JFL, SVH, and YN (single-molecule super-resolution microscopy and related image reconstructions and analysis), and RD (fluorescent imaging and image analysis). SLG, DM and CC analyzed the RNA-seq data; FM, EDM, CLW, AKG and RD participated in generating testable hypothesis, analyzing data, and reading, reviewing, and editing the manuscript. PC, JL, YN, AKG and RD wrote the manuscript.

