## Supplementary Figures for "Lysine Demethylase 4A is a Centrosome Associated Protein Required for Centrosome Integrity and Genomic Stability"

### SUPPLEMENTAL FIGURE LEGENDS

**Supplemental Figure 1. KDM4A localizes to the centrosome.**

A-C Immunoblot analysis of nuclear and cytoplasmic fractions obtained by biochemical fractionation of HEK293T (A), HKC (B) and MEFs (C) probed for the indicated antibodies.

D-E Immunoblot analysis of nuclear and cytoplasmic fractions obtained from HEK293T (D) and HKC (E) cells over expressing HA-KDM4A, probed for the indicated antibodies.

F-G Representative images from HKC cells stained for KDM4A (green) showing co-localization with γ-tubulin (red) and centrin 2 (red). Nuclei are counterstained with DAPI (blue). Scale bar – 5 μm.

H-I Co-immunoprecipitation using centrobin (H) and CP110 (I) to pull down KDM4A as indicated. Input for each sample is also shown.

**Supplemental Figure 2. Schematic of the optical setup used to collect 3D single-molecule super-resolution data.** Single fluorophores of Cy3B are excited by 560 nm using widefield epi-illumination. A 100x objective lens serves for both illumination and collection of emitted light. A 4f optical relay system is used to image the emitted light, which is split by a dichroic mirror into two emission paths. Transmissive phase masks in the Fourier plane of both paths modulate the emission light, thereby changing the shape of the point spread function (PSF) to that of the double helix PSF, which encodes the 3D position of the emitter. The two paths which are split in the 4f system are imaged onto separate regions of an EMCCD camera. Insets show the shape of the double helix PSFs in channels 1 and 2 at different axial (z) positions. Scale bar is 1 μm. The schematic is not drawn to scale.

**Supplemental Figure 3. KDM4A co-localizes with centrobin at the spindle poles.**

A-I Representative 3D super-resolution reconstructions of a spindle pole shown in the xy (A), yz (B) and xz (C) plane showing the KDM4A distribution (pseudo-colored green) intensity overlaying the distribution of the centrosome marker – centrobin (pseudo-colored magenta) during mitosis. The distributions of KDM4A (pseudo-colored green) and centrobin (pseudo-colored magenta) are also shown individually in the xy (D, G), yz (E, H) and xz (F, I) planes. Scale bar – 1 μm.

J Representative images showing immunolabelled KDM4A (green) in the four phases of mitosis (prophase, metaphase, anaphase, and telophase) in hTERT RPE-1 cells. Centrosomes are labelled with anti-centrobin antibody (red), and nuclei are counterstained with DAPI (blue). Scale bar – 5 μm.

**Supplemental Figure 4. Quantitation of the KDM4A distribution at the mitotic spindle.**

KDM4A distributions (pseudo-colored green) measured using 3D single-molecule super-resolution imaging of individual centrosomes (n=5). The three axes (xy, yz, and xz) are shown from left to right in each row with normalized histograms of the number of localizations along each axis shown below each reconstruction. Centrosome 1 (A-F), centrosome 2 (G-L), centrosome 3 (M-R), centrosome 4 (S-X), centrosome 5 (AA-FF). Scale bar – 1 μm.

**Supplemental Figure 5. KDM4A loss results in abnormal mitosis.**

A Representative image from *Kdm4a*-deficient MEFs stained for centriole marker Cep135 (green) and γ-tubulin (red). Nuclei are counterstained with DAPI (blue). Scale bar – 5 μm.

B Immunoblots of cellular lysates of *KDM4A*-proficient (siC) and deficient (siKDM4A) hTERT RPE-1 cells probed for the indicated antibodies.

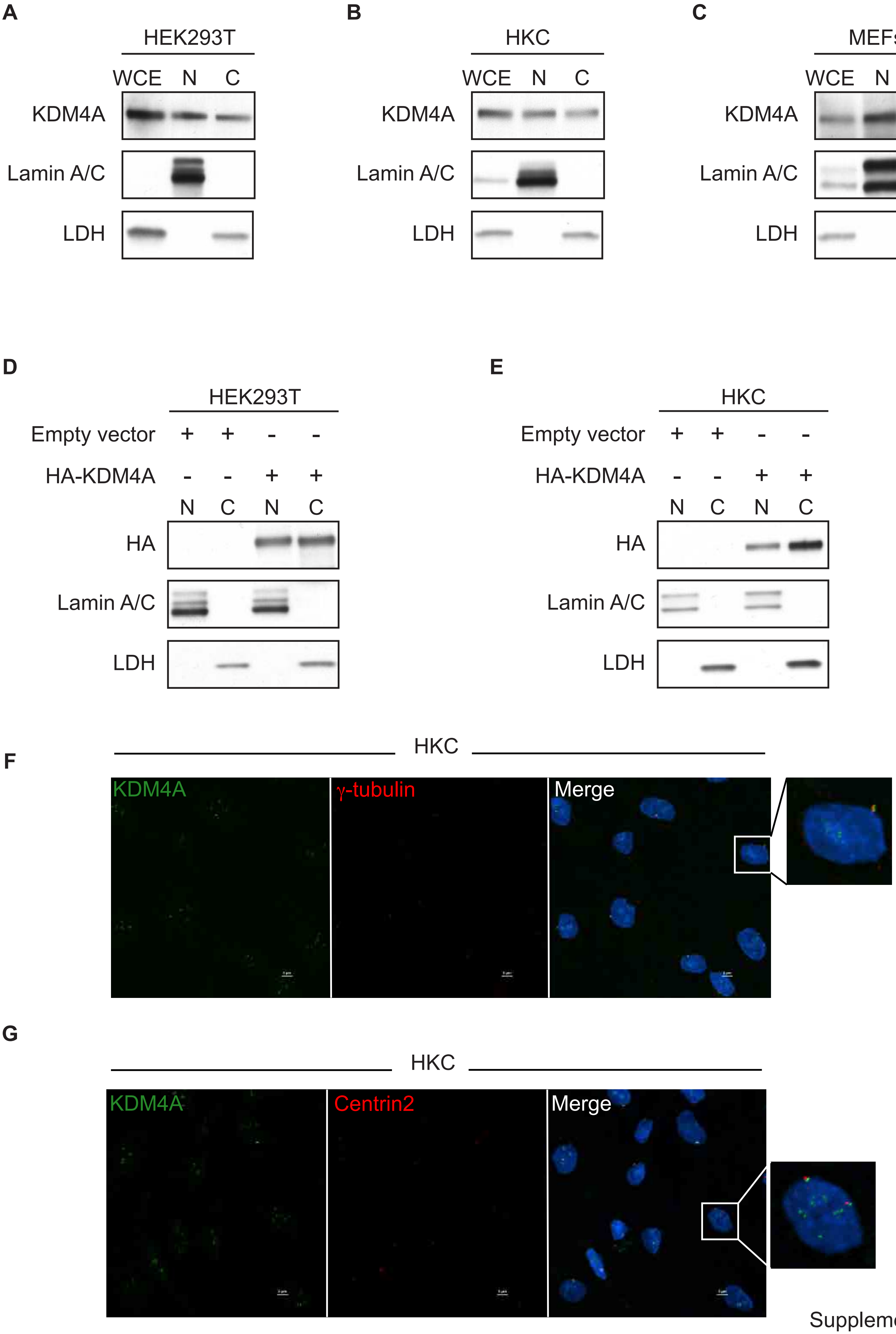

Supplemental Figure 1

H

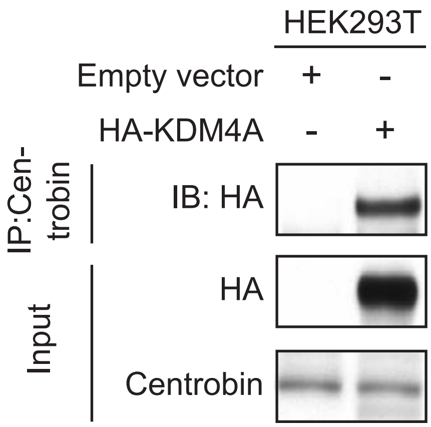
HEK293T
Empty vector + -
HA-KDM4A - +
IP:Cen-trobin
IB: HA
Input
HA
Centrobin

I

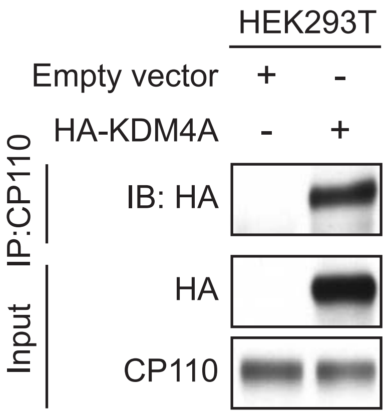
HEK293T
Empty vector + -
HA-KDM4A - +
IP:CP110
IB: HA
Input
HA
CP110

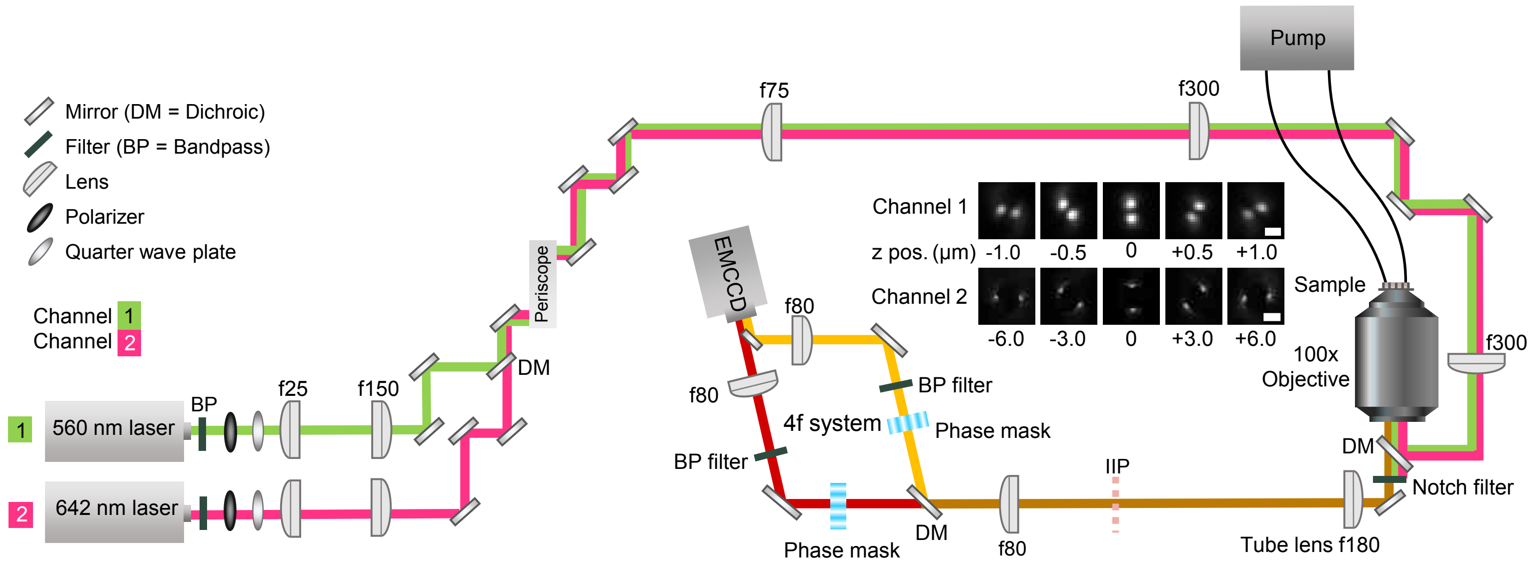

Supplemental Figure 2

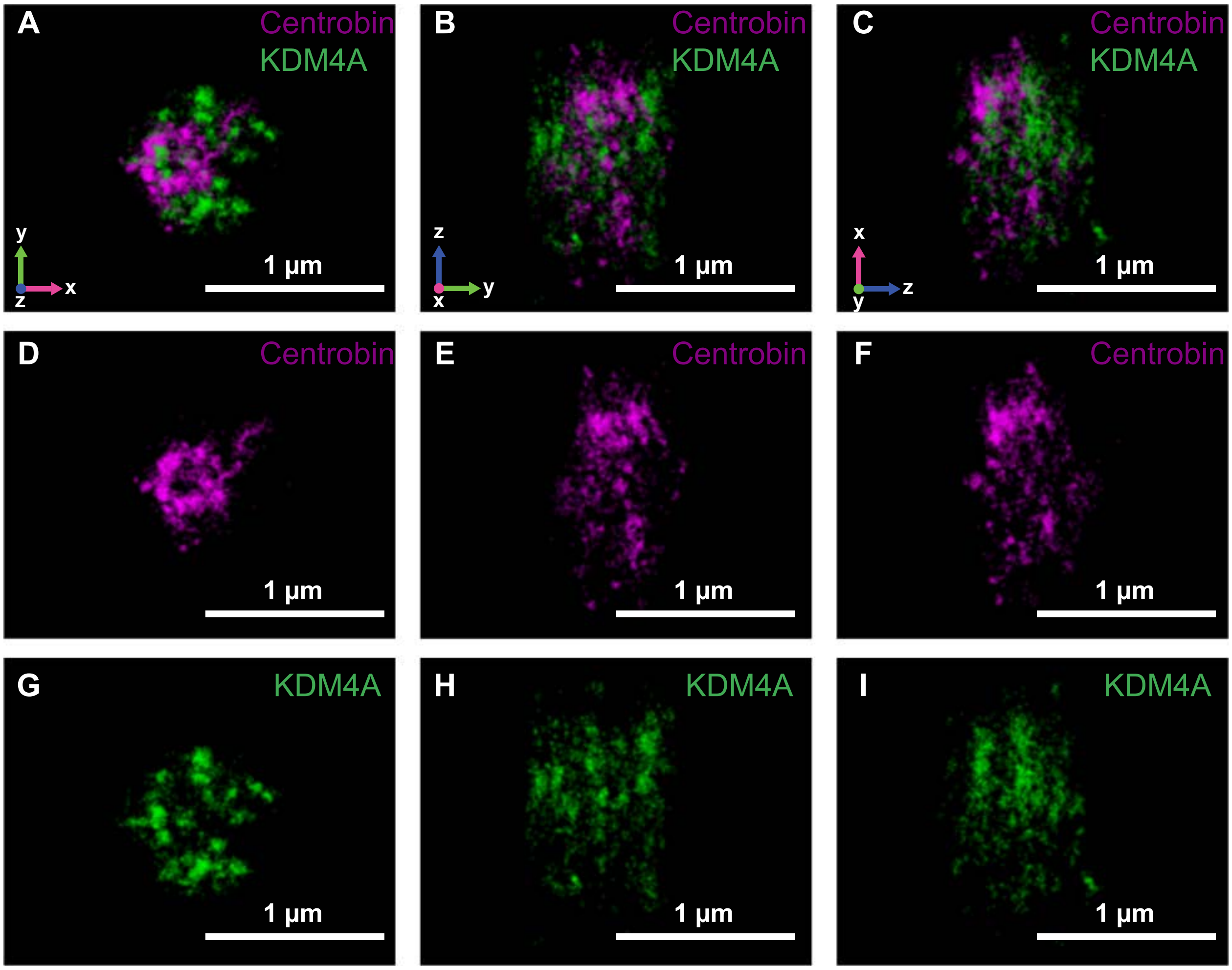

Supplemental Figure 3

**J**

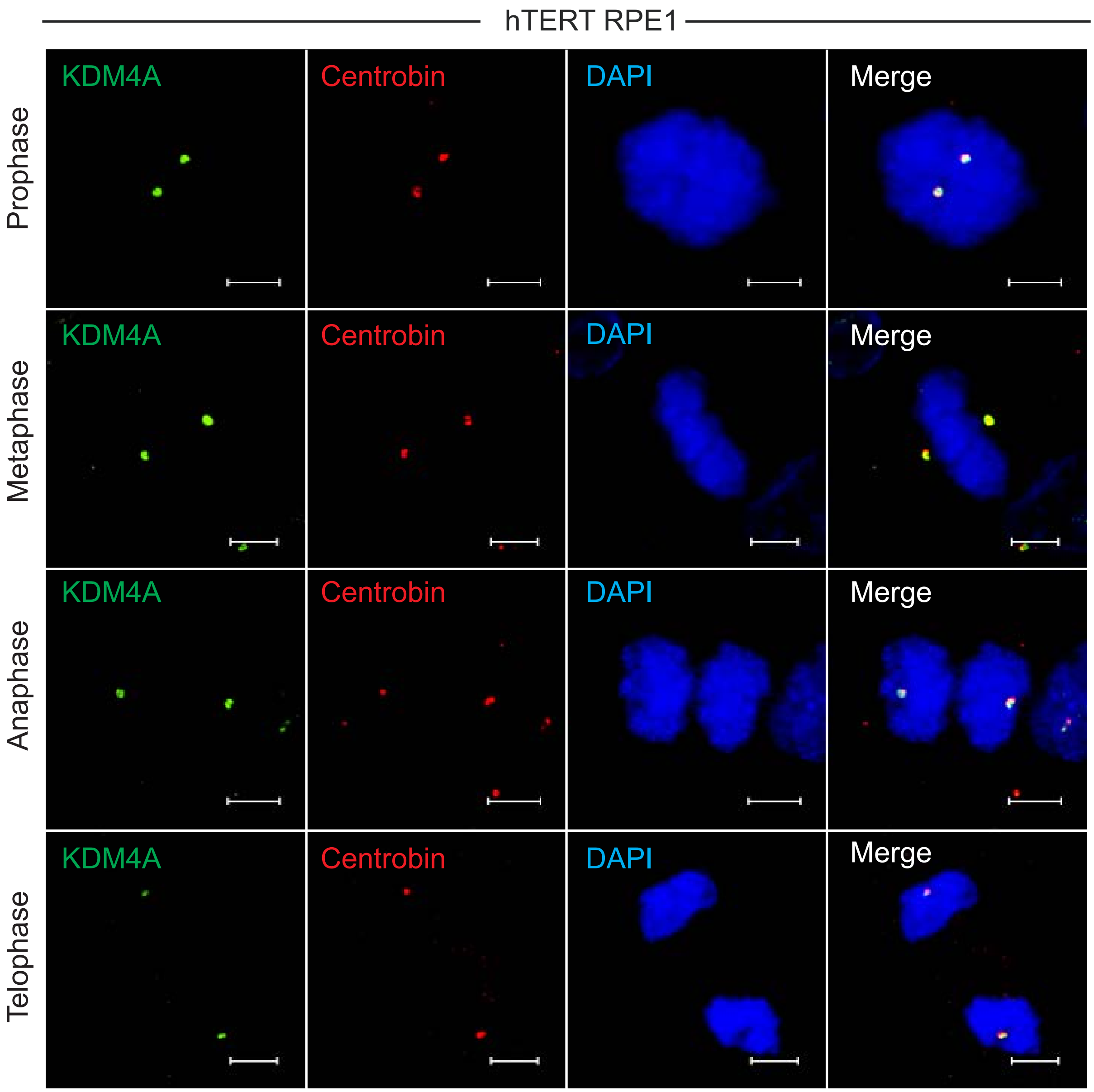
hTERT RPE1
KDM4A
Centrobin
DAPI
Merge
Prophase
Metaphase
Anaphase
Telophase

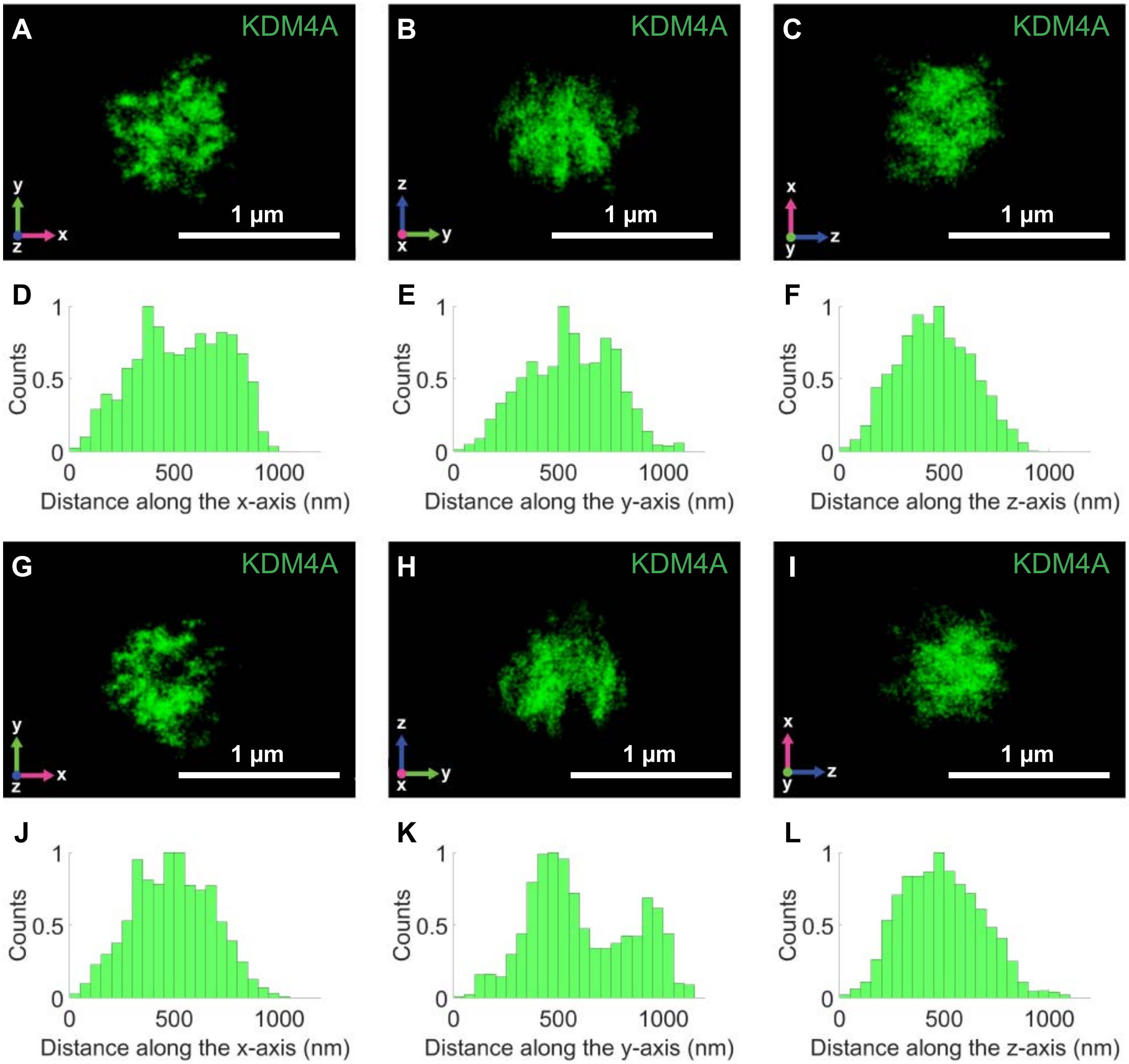

Supplemental Figure 4

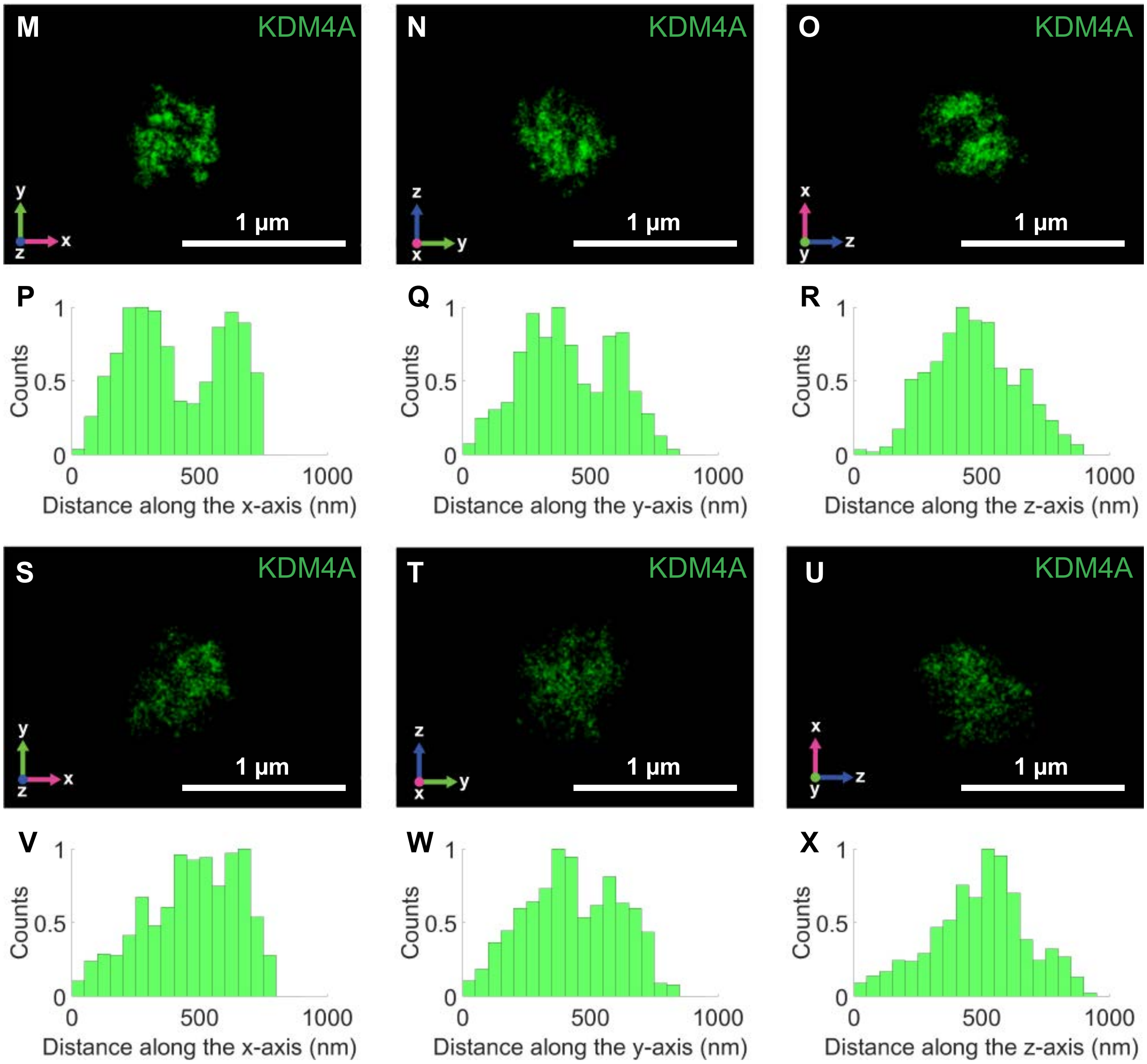
M
KDM4A
1 µm
N
KDM4A
1 µm
O
KDM4A
1 µm
P
Q
R
Counts
Distance along the x-axis (nm)
Distance along the y-axis (nm)
Distance along the z-axis (nm)
S
KDM4A
1 µm
T
KDM4A
1 µm
U
KDM4A
1 µm
V
W
X
Counts
Distance along the x-axis (nm)
Distance along the y-axis (nm)
Distance along the z-axis (nm)

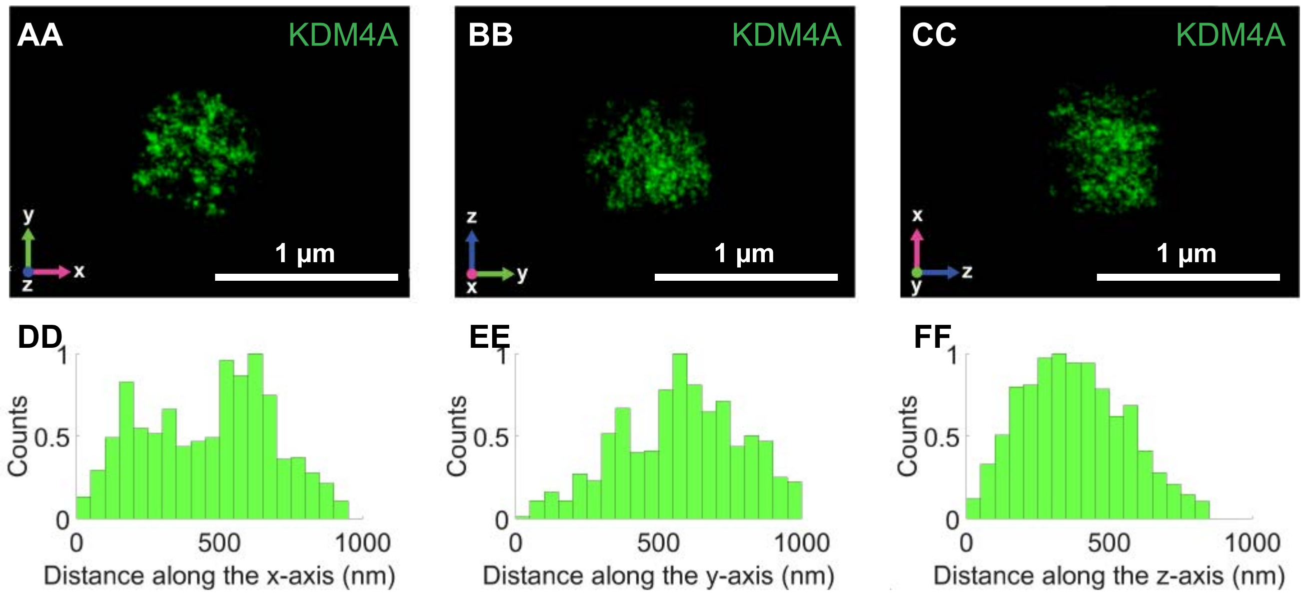
AA
KDM4A
1 µm
BB
KDM4A
1 µm
CC
KDM4A
1 µm
DD
Counts
Distance along the x-axis (nm)
EE
Counts
Distance along the y-axis (nm)
FF
Counts
Distance along the z-axis (nm)

**A**

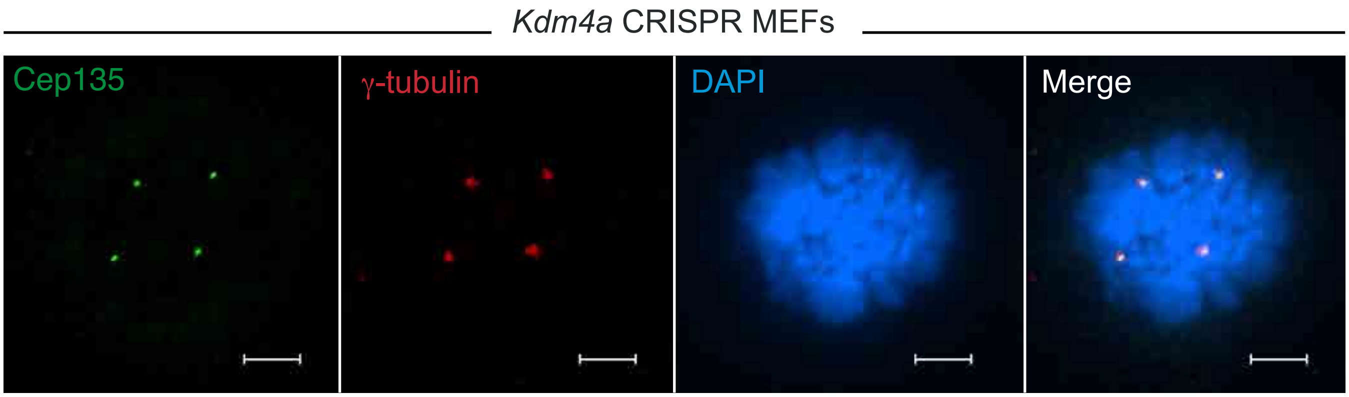
*Kdm4a* CRISPR MEFs
Cep135
γ-tubulin
DAPI
Merge

**B**

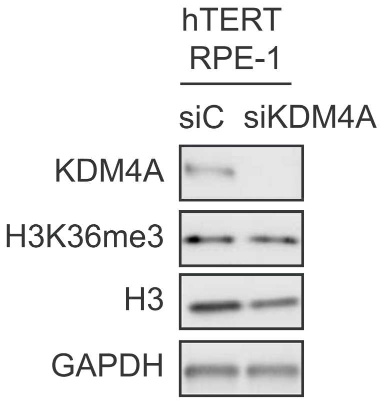
hTERT
RPE-1
siC
siKDM4A
KDM4A
H3K36me3
H3
GAPDH

| **Supplemental Table 1:** Differentially expressed genes comparing MEF NT over MEF KDM4A Crispr | | | |
|---|---|---|---|
| **GeneSymbol** | **logFC** | **PValue** | **FDR** |
| Hdx | 9.65 | 1.95329E-22 | 1.50989E-19 |
| Negr1 | 7.83 | 3.20814E-11 | 5.31406E-09 |
| Anxa8 | 7.63 | 2.40904E-10 | 3.3654E-08 |
| Cacna1b | 7.55 | 4.88176E-10 | 6.43228E-08 |
| Stag3 | 6.45 | 1.50447E-14 | 4.15342E-12 |
| Gfra2 | 6.18 | 1.16284E-34 | 2.24719E-31 |
| Cd34 | 5.90 | 1.16906E-29 | 1.50614E-26 |
| Slc24a3 | 5.81 | 1.12969E-14 | 3.27467E-12 |
| Slc16a10 | 4.59 | 2.55791E-14 | 6.89743E-12 |
| Aff2 | 4.57 | 2.85374E-11 | 4.79553E-09 |
| Hoxa9 | 4.55 | 6.69246E-10 | 8.62211E-08 |
| Sorcs2 | 4.27 | 1.56752E-20 | 9.08769E-18 |
| Bmp6 | 4.27 | 1.22787E-10 | 1.77965E-08 |
| Thy1 | 4.11 | 4.81625E-36 | 1.86148E-32 |
| Gm19410 | 4.11 | 6.50023E-08 | 5.74486E-06 |
| Nkx3-2 | 3.97 | 7.77924E-10 | 9.81925E-08 |
| Rnf128 | 3.96 | 3.05565E-15 | 9.57576E-13 |
| Blnk | 3.93 | 4.25857E-14 | 1.07344E-11 |
| Ralgps2 | 3.90 | 4.74234E-10 | 6.32039E-08 |
| Brinp3 | 3.76 | 2.43651E-09 | 2.74285E-07 |
| Fhl1 | 3.73 | 2.50677E-31 | 3.63325E-28 |
| Sgsm1 | 3.70 | 1.60805E-09 | 1.9222E-07 |
| Gpr149 | 3.70 | 7.17566E-08 | 6.20909E-06 |
| Cpeb1 | 3.64 | 5.20305E-11 | 8.37908E-09 |
| Acsbg1 | 3.58 | 9.23505E-07 | 6.18962E-05 |
| Rgs20 | 3.57 | 2.30871E-07 | 1.78831E-05 |
| Mei4 | 3.47 | 6.13796E-08 | 5.47459E-06 |
| Slc7a3 | 3.28 | 1.17745E-24 | 9.75179E-22 |
| Cystm1 | 3.27 | 7.62088E-09 | 8.03311E-07 |
| Dio2 | 3.15 | 5.05995E-08 | 4.56782E-06 |
| Adm2 | 3.03 | 7.8442E-10 | 9.81925E-08 |

| | | | |
|---|---|---|---|
| Eif4g3 | 2.96 | 5.50088E-22 | 3.98642E-19 |
| Tusc1 | 2.94 | 2.23056E-06 | 0.00013541 |
| P2rx3 | 2.90 | 2.09361E-09 | 2.42754E-07 |
| Pde8a | 2.79 | 3.90452E-16 | 1.37191E-13 |
| Ntn4 | 2.76 | 6.7968E-09 | 7.29712E-07 |
| Rasip1 | 2.75 | 7.60758E-06 | 0.000404633 |
| Gdf15 | 2.70 | 1.54969E-06 | 9.81892E-05 |
| Neto2 | 2.69 | 7.85286E-09 | 8.20306E-07 |
| Tenm4 | 2.67 | 3.41158E-11 | 5.57145E-09 |
| Neurl3 | 2.67 | 7.72162E-25 | 6.88709E-22 |
| Pld4 | 2.67 | 1.43994E-05 | 0.000704479 |
| Crybg1 | 2.66 | 4.86882E-06 | 0.000270115 |
| Lgals3 | 2.65 | 7.39553E-19 | 3.43005E-16 |
| Pla2g7 | 2.64 | 4.7313E-07 | 3.40742E-05 |
| Xkr6 | 2.63 | 1.66121E-06 | 0.000104683 |
| Tmem171 | 2.61 | 1.05205E-05 | 0.000530369 |
| Car6 | 2.59 | 3.96248E-14 | 1.021E-11 |
| Sema4f | 2.57 | 1.48866E-06 | 9.48404E-05 |
| Igdcc4 | 2.55 | 1.17418E-07 | 9.8011E-06 |
| Ccl20 | 2.54 | 8.00643E-08 | 6.87663E-06 |
| Adm | 2.50 | 1.49119E-05 | 0.000723444 |
| Spocd1 | 2.49 | 0.00014276 | 0.005221782 |
| Cox6a2 | 2.49 | 1.49777E-05 | 0.000723612 |
| Mycl | 2.44 | 8.49566E-07 | 5.76196E-05 |
| Dchs1 | 2.40 | 1.61214E-08 | 1.61145E-06 |
| Nkx2-5 | 2.31 | 3.52181E-06 | 0.000203161 |
| Klrg2 | 2.30 | 3.23918E-06 | 0.000188735 |
| Cdkn1c | 2.27 | 0.00029864 | 0.009539194 |
| Tbc1d9 | 2.26 | 8.59914E-06 | 0.000446489 |
| Uchl1 | 2.26 | 5.3235E-10 | 6.93551E-08 |
| Tspan15 | 2.24 | 5.01034E-07 | 3.56411E-05 |
| Syngr1 | 2.20 | 2.6823E-05 | 0.001229298 |
| Prrg4 | 2.13 | 5.71276E-05 | 0.002365696 |
| Il3ra | 2.10 | 1.00624E-05 | 0.000509492 |
| Samd12 | 2.10 | 0.000980488 | 0.026500589 |
| Usp17la | 2.08 | 0.000131394 | 0.004867437 |
| Als2cl | 2.08 | 5.34221E-05 | 0.002244311 |
| Mid2 | 2.07 | 2.89404E-08 | 2.72816E-06 |
| Artn | 2.06 | 4.22931E-05 | 0.001823006 |
| Hecw2 | 2.05 | 8.73783E-06 | 0.000448297 |
| Zfp951 | 2.05 | 0.000286246 | 0.009168575 |
| Tmcc3 | 2.03 | 2.87732E-06 | 0.000170217 |
| Lingo1 | 2.03 | 0.000134537 | 0.004936556 |

| | | | |
|---|---|---|---|
| Wdr66 | 2.03 | 2.75017E-05 | 0.001245635 |
| Cep126 | 2.02 | 0.000598128 | 0.017513359 |
| Tmeff2 | 1.98 | 1.35428E-05 | 0.000668207 |
| Stbd1 | 1.97 | 6.11694E-09 | 6.62859E-07 |
| Peg3 | 1.96 | 3.42979E-14 | 9.03827E-12 |
| Pik3c2b | 1.95 | 0.000512741 | 0.015402159 |
| Cdhr1 | 1.95 | 2.15026E-06 | 0.000131289 |
| C130026I21Rik | 1.94 | 0.000122948 | 0.004613552 |
| Cgn | 1.91 | 0.001069463 | 0.02827161 |
| Fzd3 | 1.90 | 0.000241528 | 0.008047457 |
| Acod1 | 1.88 | 4.92743E-09 | 5.44129E-07 |
| Msln | 1.88 | 1.84833E-09 | 2.18688E-07 |
| Adamtsl3 | 1.87 | 7.16442E-11 | 1.10762E-08 |
| Vldlr | 1.87 | 3.59194E-12 | 7.18079E-10 |
| Ankrd6 | 1.84 | 3.92263E-05 | 0.001703481 |
| Zcchc3 | 1.84 | 2.24747E-05 | 0.001042377 |
| Mettl7b | 1.80 | 0.001224516 | 0.031412077 |
| Elovl7 | 1.80 | 0.001676874 | 0.040506995 |
| Fez1 | 1.78 | 4.82794E-12 | 9.48813E-10 |
| Rab3b | 1.78 | 0.00010205 | 0.003944237 |
| Txlnb | 1.76 | 0.002117423 | 0.048140235 |
| Myom1 | 1.75 | 0.001790149 | 0.042188573 |
| Espnl | 1.74 | 0.000437001 | 0.013299288 |
| Cdh2 | 1.72 | 3.10036E-13 | 7.04875E-11 |
| Slc25a45 | 1.69 | 0.000561581 | 0.016653535 |
| Rragd | 1.68 | 5.08192E-08 | 4.56782E-06 |
| Tmsb4x | 1.67 | 1.6265E-12 | 3.36773E-10 |
| Atp1a3 | 1.67 | 2.47124E-10 | 3.41119E-08 |
| Ar | 1.66 | 5.25548E-09 | 5.74881E-07 |
| Plb1 | 1.66 | 0.000932525 | 0.025441485 |
| Nipsnap1 | 1.64 | 5.03261E-05 | 0.002129677 |
| Asxl3 | 1.63 | 0.001718311 | 0.040911331 |
| Peg12 | 1.63 | 0.00010194 | 0.003944237 |
| Hdc | 1.62 | 0.001202429 | 0.031120899 |
| Cplx2 | 1.62 | 0.000122188 | 0.004613552 |
| Glrb | 1.60 | 1.86272E-05 | 0.000888815 |
| Pdzd7 | 1.59 | 0.000220747 | 0.007550323 |
| F11r | 1.58 | 2.89393E-08 | 2.72816E-06 |
| Cbx7 | 1.58 | 0.000261577 | 0.008543628 |
| Itgb7 | 1.58 | 1.5975E-11 | 2.80652E-09 |
| Pdlim1 | 1.58 | 2.29294E-07 | 1.78831E-05 |
| Reep6 | 1.51 | 1.48581E-07 | 1.18813E-05 |
| Dtna | 1.50 | 5.36956E-06 | 0.000293679 |

| | | | |
|---|---|---|---|
| Fbxl16 | 1.50 | 0.000339589 | 0.010699817 |
| Cenpv | 1.48 | 4.16089E-05 | 0.001800205 |
| Gm4070 | 1.46 | 1.33629E-06 | 8.60791E-05 |
| Cd24a | 1.46 | 2.25355E-09 | 2.58712E-07 |
| Ptpn22 | 1.45 | 0.001817876 | 0.04266857 |
| Dhrs3 | 1.44 | 3.79013E-06 | 0.000216486 |
| Strbp | 1.44 | 8.28235E-06 | 0.000434542 |
| Mfsd2a | 1.43 | 0.000591107 | 0.017351608 |
| Stau2 | 1.43 | 2.75122E-06 | 0.000163592 |
| Adam22 | 1.43 | 0.001452683 | 0.036223342 |
| Plekhh1 | 1.42 | 0.001115059 | 0.02925137 |
| Slc4a4 | 1.41 | 4.79281E-07 | 3.43041E-05 |
| AA467197 | 1.40 | 8.54728E-07 | 5.76196E-05 |
| Id1 | 1.40 | 5.18086E-06 | 0.000286058 |
| Gm14137 | 1.40 | 6.21375E-05 | 0.002545881 |
| Hspb1 | 1.40 | 2.30397E-09 | 2.61908E-07 |
| Tuft1 | 1.39 | 0.000845368 | 0.023617709 |
| Mogat2 | 1.39 | 0.000585017 | 0.017216431 |
| Ndrg2 | 1.38 | 3.0175E-05 | 0.0013614 |
| Nxph4 | 1.38 | 0.001471492 | 0.03661363 |
| 4930452B06Rik | 1.35 | 0.000255788 | 0.008425746 |
| Fer1l5 | 1.33 | 0.000132135 | 0.004879302 |
| Man1c1 | 1.30 | 1.06305E-05 | 0.000533597 |
| Lamc2 | 1.30 | 0.001217879 | 0.031311092 |
| Timp3 | 1.28 | 4.4383E-08 | 4.05213E-06 |
| Ereg | 1.27 | 4.27734E-08 | 3.93617E-06 |
| Angptl6 | 1.25 | 0.001090661 | 0.028676213 |
| Nqo1 | 1.24 | 0.000385911 | 0.011932364 |
| Rnd1 | 1.24 | 1.17495E-07 | 9.8011E-06 |
| Gvin1 | 1.24 | 2.05297E-05 | 0.000968912 |
| Paqr3 | 1.23 | 0.000102928 | 0.003964946 |
| Itga6 | 1.23 | 2.89166E-07 | 2.20584E-05 |
| Hfe | 1.23 | 0.000158356 | 0.005684622 |
| Nos2 | 1.21 | 3.85232E-06 | 0.000217891 |
| Ifi206 | 1.21 | 8.34349E-05 | 0.003298795 |
| Enpp4 | 1.20 | 0.00021575 | 0.007423201 |
| Cd74 | 1.20 | 3.11604E-06 | 0.000182477 |
| Sema3c | 1.18 | 4.31802E-07 | 3.18901E-05 |
| Sema6b | 1.17 | 0.001548148 | 0.038112046 |
| Pim1 | 1.16 | 0.000145287 | 0.005280881 |
| Ripk3 | 1.16 | 8.52352E-07 | 5.76196E-05 |
| Dnaja4 | 1.15 | 0.000878552 | 0.024370367 |
| Chd7 | 1.15 | 0.00037258 | 0.011613077 |

| | | | |
|---|---|---|---|
| Shc2 | 1.14 | 6.90928E-05 | 0.002791396 |
| Mreg | 1.12 | 0.001984279 | 0.046107635 |
| Bok | 1.11 | 0.000258723 | 0.008498266 |
| Pxdc1 | 1.10 | 0.000536799 | 0.016083156 |
| Gfra1 | 1.10 | 0.000304194 | 0.0096899 |
| Klf4 | 1.10 | 2.15134E-06 | 0.000131289 |
| Cdkn1a | 1.10 | 2.57657E-06 | 0.000154795 |
| H2-M3 | 1.09 | 0.000228116 | 0.007711389 |
| Cdsn | 1.09 | 2.73084E-06 | 0.000163217 |
| Gprc5a | 1.09 | 3.53619E-05 | 0.001564965 |
| Ero1l | 1.07 | 6.03582E-05 | 0.002490581 |
| Alcam | 1.06 | 0.00046026 | 0.013897708 |
| Cth | 1.06 | 5.30005E-06 | 0.000291252 |
| Lcn2 | 1.06 | 0.000129741 | 0.004837121 |
| Epb41 | 1.05 | 1.45336E-05 | 0.000708055 |
| Slpi | 1.05 | 9.9049E-06 | 0.000503716 |
| Hbegf | 1.04 | 6.1064E-06 | 0.000330858 |
| Slc38a1 | 1.04 | 6.62361E-06 | 0.000353921 |
| Sox12 | 1.03 | 8.43681E-06 | 0.000440652 |
| Magi1 | 1.03 | 0.000240345 | 0.008031139 |
| Plk2 | 1.02 | 8.29775E-05 | 0.003294947 |
| Lgi2 | 1.01 | 0.000459483 | 0.013897708 |
| Zfp469 | -1.00 | 2.05565E-05 | 0.000968912 |
| Sbk1 | -1.00 | 0.001316093 | 0.033174115 |
| Lbp | -1.02 | 8.63099E-05 | 0.003392418 |
| Slc1a3 | -1.03 | 8.19811E-06 | 0.000432078 |
| Mfsd13a | -1.03 | 0.000231748 | 0.007811377 |
| Zfp874a | -1.05 | 0.000580874 | 0.017137988 |
| Mmp9 | -1.05 | 8.01476E-06 | 0.000424343 |
| Gm8113 | -1.05 | 0.000189572 | 0.00670149 |
| Cxcl5 | -1.06 | 0.000111094 | 0.004264801 |
| Evi2a | -1.07 | 0.001897568 | 0.044359469 |
| Pcdhb19 | -1.08 | 0.000993536 | 0.026666779 |
| Rab3il1 | -1.08 | 1.97283E-05 | 0.000937497 |
| Ada | -1.08 | 0.001996311 | 0.04620204 |
| D130040H23Rik | -1.09 | 0.002109443 | 0.048053032 |
| Sh3rf3 | -1.10 | 0.000114386 | 0.004362852 |
| Tnxb | -1.11 | 0.000382178 | 0.011880318 |
| Tnip3 | -1.11 | 0.000308686 | 0.009806064 |
| Ppm1h | -1.12 | 2.01939E-06 | 0.000124547 |
| Rnf150 | -1.15 | 6.27676E-06 | 0.00033694 |
| Enpp1 | -1.16 | 1.70642E-06 | 0.000106951 |
| Psmb9 | -1.16 | 0.001079224 | 0.028439996 |

| | | | |
|---|---|---|---|
| Gm4951 | -1.16 | 1.11042E-05 | 0.000554969 |
| Arhgap31 | -1.16 | 4.55705E-07 | 3.31216E-05 |
| Pcdhb20 | -1.17 | 0.000931973 | 0.025441485 |
| Inhbb | -1.17 | 1.95501E-06 | 0.00012168 |
| Abi3bp | -1.18 | 4.57046E-07 | 3.31216E-05 |
| Ldhb | -1.18 | 8.70391E-06 | 0.000448297 |
| Clec2d | -1.18 | 2.74413E-05 | 0.001245635 |
| Ndnf | -1.19 | 3.42159E-07 | 2.55957E-05 |
| Kng2 | -1.19 | 3.08879E-07 | 2.32562E-05 |
| Mme | -1.19 | 2.96139E-07 | 2.24427E-05 |
| Eya1 | -1.19 | 0.001300816 | 0.032932226 |
| Lef1 | -1.21 | 3.59771E-05 | 0.001586138 |
| Mpeg1 | -1.21 | 0.001674682 | 0.040506995 |
| Col5a3 | -1.21 | 1.80681E-07 | 1.42517E-05 |
| Col11a1 | -1.22 | 1.38109E-07 | 1.12158E-05 |
| Maf | -1.22 | 1.36389E-07 | 1.12158E-05 |
| Gpr162 | -1.22 | 0.000649509 | 0.018827629 |
| Sulf2 | -1.23 | 9.58786E-06 | 0.000489741 |
| Pcdhb17 | -1.24 | 3.71061E-05 | 0.00162012 |
| Pcdh7 | -1.24 | 1.31413E-07 | 1.08838E-05 |
| Elfn1 | -1.25 | 9.27041E-08 | 7.90371E-06 |
| Tmtc4 | -1.26 | 6.58962E-08 | 5.74486E-06 |
| Pde1a | -1.28 | 3.80795E-08 | 3.53226E-06 |
| Hdac9 | -1.28 | 0.001032484 | 0.02745793 |
| Il1rn | -1.29 | 2.42436E-07 | 1.86162E-05 |
| Ifi27l2a | -1.29 | 5.69766E-06 | 0.000310161 |
| Plekhg4 | -1.29 | 6.34194E-05 | 0.002589251 |
| Tnc | -1.30 | 1.38323E-07 | 1.12158E-05 |
| Sox5 | -1.30 | 3.39557E-05 | 0.001514293 |
| Ptn | -1.31 | 2.88214E-08 | 2.72816E-06 |
| Cacnb2 | -1.31 | 0.0004206 | 0.012901749 |
| Hsd11b1 | -1.33 | 3.27445E-06 | 0.000189836 |
| Cthrc1 | -1.33 | 1.24219E-08 | 1.26344E-06 |
| Mgp | -1.34 | 1.21353E-06 | 7.90498E-05 |
| Pitx2 | -1.35 | 1.56575E-07 | 1.24348E-05 |
| Itga10 | -1.36 | 3.48983E-05 | 0.001550368 |
| Wnt5b | -1.37 | 0.001289583 | 0.032763204 |
| Serpina3i | -1.37 | 0.000204116 | 0.007107272 |
| F830016B08Rik | -1.38 | 6.20432E-06 | 0.0003346 |
| Thbs2 | -1.39 | 7.51773E-09 | 7.99707E-07 |
| Prkg1 | -1.40 | 4.08532E-06 | 0.000229948 |
| Gdpd2 | -1.40 | 0.001344773 | 0.033750305 |
| Creb3l1 | -1.42 | 9.60525E-09 | 9.94401E-07 |

| | | | |
|---|---|---|---|
| Igsf10 | -1.42 | 2.73994E-08 | 2.64746E-06 |
| Clip4 | -1.43 | 5.65994E-07 | 3.95343E-05 |
| P2ry10b | -1.44 | 0.000211893 | 0.007312191 |
| Tnfrsf11b | -1.44 | 3.77103E-07 | 2.80289E-05 |
| Cxcl2 | -1.45 | 4.94551E-05 | 0.002100483 |
| Cpxm1 | -1.48 | 4.4816E-07 | 3.28887E-05 |
| Angpt4 | -1.48 | 1.45981E-07 | 1.17545E-05 |
| Cyp7b1 | -1.49 | 2.87561E-10 | 3.92266E-08 |
| Gm5617 | -1.50 | 0.002085352 | 0.047789241 |
| Aqp1 | -1.52 | 5.22784E-07 | 3.69615E-05 |
| Gda | -1.52 | 0.001592808 | 0.039045687 |
| Map3k5 | -1.52 | 1.39005E-06 | 8.90476E-05 |
| Ppbp | -1.52 | 4.0666E-09 | 4.53387E-07 |
| Ppfibp2 | -1.53 | 1.41298E-05 | 0.000694218 |
| Fgfr2 | -1.53 | 1.42267E-10 | 2.03653E-08 |
| Abca9 | -1.54 | 8.26386E-11 | 1.26078E-08 |
| Mctp2 | -1.54 | 0.000364719 | 0.011398709 |
| Islr | -1.56 | 1.02892E-10 | 1.52953E-08 |
| Prr33 | -1.56 | 0.001822264 | 0.042685143 |
| Kng1 | -1.57 | 1.81411E-11 | 3.13949E-09 |
| Sned1 | -1.57 | 5.47773E-11 | 8.70058E-09 |
| Fat4 | -1.58 | 1.56256E-11 | 2.78736E-09 |
| F8 | -1.59 | 0.001332277 | 0.033509211 |
| Hsh2d | -1.60 | 0.000545346 | 0.016297134 |
| Ntrk3 | -1.60 | 0.000275634 | 0.008927293 |
| Hivep3 | -1.61 | 2.28143E-08 | 2.24179E-06 |
| Csgalnact1 | -1.63 | 0.000249354 | 0.008268982 |
| Tspan13 | -1.65 | 0.000995861 | 0.026667449 |
| Olfml3 | -1.65 | 2.55684E-11 | 4.35978E-09 |
| Cftr | -1.65 | 0.001691283 | 0.040601287 |
| Car9 | -1.66 | 0.00067425 | 0.019448158 |
| Ccn5 | -1.67 | 1.46356E-12 | 3.08544E-10 |
| Ccl9 | -1.71 | 5.10859E-13 | 1.09693E-10 |
| Tgfbi | -1.72 | 2.24513E-12 | 4.56706E-10 |
| Il16 | -1.73 | 5.48266E-05 | 0.002286744 |
| Col23a1 | -1.74 | 5.32851E-12 | 1.02973E-09 |
| Prr16 | -1.74 | 2.27365E-05 | 0.001050318 |
| Kcnj15 | -1.74 | 1.82203E-10 | 2.5764E-08 |
| Serpinb1a | -1.74 | 1.31623E-08 | 1.3271E-06 |
| Igfbp2 | -1.74 | 9.5527E-12 | 1.73068E-09 |
| Batf2 | -1.77 | 0.001769339 | 0.041838976 |
| Tgtp2 | -1.78 | 1.59355E-09 | 1.9222E-07 |
| Angptl8 | -1.80 | 1.13611E-07 | 9.61551E-06 |

| | | | |
|---|---|---|---|
| Apba1 | -1.80 | 0.00067427 | 0.019448158 |
| Stra6 | -1.80 | 1.1092E-08 | 1.13816E-06 |
| Slco1a5 | -1.81 | 4.66001E-13 | 1.03909E-10 |
| Phactr1 | -1.85 | 0.00015763 | 0.005676145 |
| Tgfbr3l | -1.85 | 0.000342622 | 0.01076613 |
| Cxcl3 | -1.86 | 7.87572E-10 | 9.81925E-08 |
| Ism1 | -1.89 | 2.92977E-08 | 2.73957E-06 |
| Mertk | -1.90 | 1.00257E-06 | 6.64271E-05 |
| Enpp3 | -1.90 | 0.00073717 | 0.021060139 |
| Peli2 | -1.91 | 0.000988936 | 0.026604898 |
| Arhgap8 | -1.91 | 0.000491525 | 0.014803189 |
| Cxcl12 | -1.93 | 7.31399E-14 | 1.80438E-11 |
| Shroom3 | -1.95 | 0.000191832 | 0.00676078 |
| Chst15 | -1.97 | 1.42724E-16 | 5.1715E-14 |
| Col6a3 | -1.98 | 4.96662E-13 | 1.08656E-10 |
| 4930578I06Rik | -1.98 | 0.001619555 | 0.039617593 |
| Akap12 | -1.99 | 1.41795E-14 | 4.01004E-12 |
| Lifr | -2.00 | 2.85418E-15 | 9.19283E-13 |
| Nat8f4 | -2.05 | 4.37057E-06 | 0.000244815 |
| Ogn | -2.05 | 7.54816E-18 | 3.12575E-15 |
| Ms4a4d | -2.06 | 8.0839E-18 | 3.23217E-15 |
| Mcoln2 | -2.09 | 3.81348E-06 | 0.000216751 |
| Aldh1l1 | -2.11 | 2.50435E-15 | 8.29656E-13 |
| Sntb1 | -2.12 | 1.10919E-13 | 2.6247E-11 |
| Tbx2 | -2.12 | 1.75157E-05 | 0.000839234 |
| Csf2rb | -2.13 | 1.12604E-05 | 0.000560362 |
| Lrrc4c | -2.13 | 0.001184039 | 0.030713495 |
| Slitrk6 | -2.14 | 8.07942E-12 | 1.487E-09 |
| Plxdc1 | -2.14 | 2.31347E-07 | 1.78831E-05 |
| Igf1 | -2.15 | 2.97573E-06 | 0.000175145 |
| Dpt | -2.15 | 2.5291E-06 | 0.000152734 |
| Daam2 | -2.17 | 1.96241E-06 | 0.00012168 |
| Tro | -2.19 | 0.000227515 | 0.007711389 |
| Agrp | -2.20 | 0.000770284 | 0.021837265 |
| Omd | -2.20 | 7.30921E-05 | 0.002932537 |
| Mmp10 | -2.20 | 1.32137E-05 | 0.000654757 |
| Nat8f1 | -2.22 | 1.31264E-09 | 1.60211E-07 |
| Tmem246 | -2.24 | 3.09134E-05 | 0.001389305 |
| Serpina3g | -2.24 | 1.47429E-17 | 5.51431E-15 |
| Gm5662 | -2.27 | 3.16911E-10 | 4.27277E-08 |
| Adamts9 | -2.28 | 1.41279E-18 | 6.30052E-16 |
| Kbtbd11 | -2.30 | 1.12358E-06 | 7.40221E-05 |
| Crocc2 | -2.32 | 1.25914E-17 | 4.86659E-15 |

| | | | |
|---|---|---|---|
| Sp5 | -2.33 | 1.18178E-06 | 7.74163E-05 |
| Mfap5 | -2.33 | 6.12892E-05 | 0.00252003 |
| Robo2 | -2.36 | 2.28482E-20 | 1.26155E-17 |
| Lum | -2.39 | 9.29175E-21 | 5.67041E-18 |
| Rp1l1 | -2.40 | 2.58828E-05 | 0.001190918 |
| Batf | -2.41 | 5.71271E-05 | 0.002365696 |
| Dlgap2 | -2.42 | 9.77943E-07 | 6.51681E-05 |
| Nrep | -2.43 | 3.7018E-21 | 2.38458E-18 |
| Madcam1 | -2.47 | 5.44126E-05 | 0.002277669 |
| Slco1a6 | -2.47 | 6.38578E-11 | 1.00058E-08 |
| Col14a1 | -2.47 | 1.26893E-06 | 8.21968E-05 |
| Aspg | -2.49 | 0.000145097 | 0.005280881 |
| Tenm3 | -2.54 | 5.5386E-25 | 5.35168E-22 |
| Serpinb2 | -2.55 | 1.96929E-25 | 2.07581E-22 |
| Itm2a | -2.58 | 9.96747E-20 | 5.0249E-17 |
| Gm10226 | -2.59 | 7.91112E-05 | 0.003152215 |
| Serpina3f | -2.67 | 2.61739E-18 | 1.12402E-15 |
| B4galnt3 | -2.68 | 4.78618E-06 | 0.000266807 |
| Postn | -2.69 | 1.08519E-19 | 5.24284E-17 |
| Ehf | -2.71 | 1.0454E-10 | 1.53436E-08 |
| Grin2a | -2.71 | 6.41837E-07 | 4.42982E-05 |
| Igf2 | -2.73 | 8.29698E-15 | 2.46676E-12 |
| Adam12 | -2.78 | 7.68509E-07 | 5.2727E-05 |
| Nr5a2 | -2.78 | 3.7167E-05 | 0.00162012 |
| Nat8 | -2.81 | 8.73383E-10 | 1.07733E-07 |
| Scara5 | -2.86 | 2.0154E-09 | 2.36046E-07 |
| Tnnt3 | -2.86 | 8.62558E-06 | 0.000446489 |
| Bhlhe22 | -2.89 | 5.2622E-07 | 3.69789E-05 |
| Stc1 | -2.90 | 1.40966E-31 | 2.33501E-28 |
| Pgm5 | -2.98 | 7.59181E-14 | 1.8339E-11 |
| Aoc3 | -3.02 | 3.10371E-05 | 0.001389478 |
| Gys2 | -3.07 | 3.73965E-06 | 0.00021466 |
| Nat8f3 | -3.09 | 6.58481E-08 | 5.74486E-06 |
| Rnf165 | -3.12 | 1.71317E-08 | 1.6978E-06 |
| Mmp3 | -3.23 | 8.07315E-35 | 1.87216E-31 |
| Mmp8 | -3.49 | 1.17314E-21 | 8.00147E-19 |
| Gad1 | -3.66 | 2.94554E-35 | 8.53839E-32 |
| Hunk | -3.78 | 6.81603E-28 | 7.90318E-25 |
| Svep1 | -3.81 | 8.61484E-20 | 4.54041E-17 |
| C1qtnf7 | -3.90 | 9.55986E-11 | 1.43957E-08 |
| Col25a1 | -3.92 | 6.25854E-07 | 4.34537E-05 |
| Col15a1 | -4.44 | 1.97466E-39 | 1.14481E-35 |
| Ednra | -4.46 | 8.10866E-16 | 2.76529E-13 |

| | | | |
|---|---|---|---|
| Sfrp1 | -4.50 | 8.5692E-58 | 9.93598E-54 |
| Mafb | -4.54 | 2.31076E-08 | 2.25154E-06 |
| Alpl | -4.78 | 6.87366E-12 | 1.30656E-09 |
| Tmem252 | -4.97 | 7.4269E-12 | 1.38895E-09 |
| Nat8f5 | -5.02 | 7.40111E-15 | 2.25831E-12 |
| Aldh1a7 | -5.21 | 0.000268927 | 0.008759011 |
| Kank4 | -8.31 | 1.91204E-13 | 4.43403E-11 |
